# A general model for two-locus assortative mating analysis and prediction

**DOI:** 10.1101/2024.01.20.576388

**Authors:** Reginald D. Smith

## Abstract

The analytical framework to investigate the effect of assortative mating for phenotypes that are determined by two loci is over a hundred years old and well established by Jennings, Wentworth, Remick, Robbins, and Wright. However, some known aspects of assortative mating have not received a concise analytical analysis or clear expression. Chief among these are linkage disequilibrium and identity disequilibrium but also the change in heterozygosity outside of the simple case of *P*_*A*_ = *P*_*B*_ = 1*/*2. In order to understand the effects of assortative mating in more detail, a general model is proposed that uses recursive difference equations rather than simulation to solve most assortative mating problems. Using this, we will expand the cases for which an exact or approximate expression for population genetic variables can be determined. In particular, the first closed form expressions for linkage disequilibrium and identity disequilibrium for two-locus assortative mating are given. We also show that assortative mating does not generate linkage disequilibrium when the loci are completely linked. Finally, this model will be used to investigate two possible cases of two-locus assortative mating in human populations regarding variants for non-syndromic deafness and the variants that largely determine blue eye color.

## 1. Introduction

Assortative mating is the propensity for mating based on phenotypic resemblance in a population. Even though phenotypically similar mates can occur in random mating, when assortative mating is present, mates are much more likely to have a phenotypic resemblance than if mating amongst phenotypes are random. Its opposite, disassortative mating, is a propensity for mates to not be phenotypically similar. While the resemblance of husbands, wives, or partners has probably long been known, the modern study of assortative mating began when Karl Pearson collected and began to quantitatively analyze tables of quantitative and categorical characters for spouses originally recorded by Francis Galton. Among other characters, he described the correlation between husbands and wives for stature and eye color (Pearson, 1900). However, since Pearson was a proponent of the biometric school, the theoretical analysis of the effects of assortative mating on phenotypes with underlying Mendelian genetic inheritance was pioneered by American geneticists including Herbert Spencer Jennings (Jennings, 1916, 1917), Wentworth & Remick (Wentworth & Remick, 1916), Rainard Robbins (Robbins, 1918), and Sewall Wright (Wright, 1921). The findings of reduced heterozygosity as well as linkage disequilibrium was established by their works. Fisher addressed assortative mating of continuous characters in his celebrated 1918 paper combining the insights of biometry and Mendelian inheritance to establish quantitative genetics (Fisher, 1918).

The next substantial theoretical advance on one and two-locus assortative mating was in (Crow & Felsenstein, 1968) who established many key results for the equilibrium genotype frequencies and genetic variance due to assortative mating at a single locus. In an exposition article (Lewtonin et. al., 1968) on non-random mating and its key effects Lewontin, Kirk, and Crow made the following distinctions:

Selective mating is character specific and involves gene frequency change; assortative mating is character specific but involves no gene frequency change; inbreeding is not character specific and involves no gene frequency change.

This statement was slightly incorrect as pointed out by (Spencer, 1992) in that it draws too distinct a line between assortative mating and selective mating. One key insight was that while assortative mating maintained gene frequencies, disassortative mating often required that gene frequencies change due to the frequency mismatches between phenotype groups. This will be expanded on later in the paper where we show Spencer is correct for most cases of disassortative mating. Further work in Ghai (1973) established the equilibrium frequencies of two-locus double homozygous genotypes for arbitrary values of loci allele frequencies when the assortative mating correlation within phenotype groups is 1.

The latest substantial theoretical work on two-locus assortative mating was given by (Hedrick, 2017). He revisited many of the past results as well as investigated assortative mating in simulations using a new parameter, *A*, to measure assortative mating by proportion of within group mating. His focus was determining linkage disequilibrium in assortative mating, a poorly covered topic in the past. He mostly uses simulation to analyze linkage disequilibrium and two-locus genotype frequencies for many conditions of allele frequencies and assortative mating at two loci. He also raised the point that recombination itself does not change the equilibrium value of linkage disequilibrium but slows its increase per generation.

### 1.1 Wright’s 1921 analysis

Probably the most influential paper on assortative mating was that of Sewall Wright in 1921 (Wright, 1921). In this paper, he described a model of assortative mating where there were five separate phenotype groups with one to three genotypes in each (See Table 1). In the assortative mating example, each phenotype group had a correlation *m* of mating with another individual from the same phenotype group and an equal and opposite correlation to mating with other phenotype groups. The most analytically tractable situation was that of the two loci being equal at 1*/*2 so that *P*_*A*_ = *P*_*B*_ = 1*/*2. Wright first showed how the heterozygosity decreased in each generation and then using the method of path coefficients calculated the expected heterozygosity after a large number of generations of assortative mating. He was able to demonstrate this for an arbitrary number of loci (his equations on the top of page 153). For two loci, the final heterozygosity could be re-written as

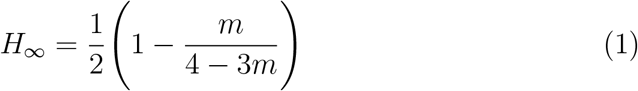

Then when *m* = 0, heterozygosity is 1*/*2 as expected and when *m* = 1 heterozygosity disappears. The expression *m/*(4 *−* 3*m*) is the effective fixation index of the population

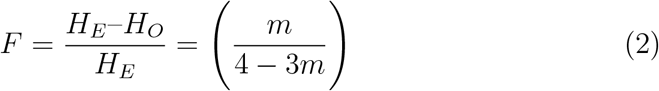

where *H*_*E*_ is the expected heterozygosity at Hardy-Weinberg equilibrium and *H*_*O*_ is the observed heterozygosity. Wright addresses linkage disequilibrium as the correlation between sister gametes, however, while he does show that the equilibrium correlation between alleles is 1 for assortative mating of *m* = 1 (corresponding to *D* = 0.25), he does not present a comprehensive expression for the equilibrium correlation for other values of *m*. Wright also excludes any analysis of the impact of recombination, though this was addressed earlier in (Robbins, 1918).

**Table 1:** The phenotype groups discussed in (Wright, 1921) and (Crow & Kimura, 1970). The double heterozygotes in phenotype group 2 are separated based on the two pairs of gamete combinations. Under assortative mating, mating is only between genotypes in the same phenotype group while in Wright’s disassortative mating model matings occur between groups as 0 *×* 4, 1 *×* 3 or 2 *×* 2.

| Phenotype Group | Genotypes |
| --- | --- |
| 0 | *aabb* |
| 1 | *aaBb*, *Aabb* |
| 2 | *AAbb*, *aaBB*, *AB/ab*, *Ab/aB* |
| 3 | *AABb*, *AaBB* |
| 4 | *AABB* |

Wright briefly addressed disassortative mating showing that the heterozygosity and fixation index can be predicted using negative values of *m* where *P*_*A*_ = *P*_*B*_ = 1*/*2.

## 2. Linkage disequilibrium, recombination, and identity disequilibrium

While many papers since Wright’s analysis have discussed linkage disequilibrium, a general formula outside of the well-known case of *P*_*A*_ = *P*_*B*_ = 1*/*2 and *m* = 1 has not yet been presented. Also, outside of (Robbins, 1918; Hedrick, 2017), the impact of recombination on the value of the final linkage disequilibrium is rarely addressed.

In addition, the question of the identity disequilibrium (Haldane, 1949; Cockerham & Weir, 1973; Smith, 2023), the correlation between heterozygotes seen across two loci when inbreeding is present, has not been comprehensively addressed since being first brought up and analyzed numerically in (Hedrick, 2017).

In order to answer these questions, and many others, it helps to have a comprehensive and flexible analytical framework to use on a variety of assortative mating models. So far equilibrium calculations in simple cases and computer based simulations have been the main tools to analyze assortative mating. This paper will introduce a general analytical framework that can be applied to create sets of recursive difference equations that can be used to analyze assortative mating and its equilibrium parameter values.

The advantage of this method over numerical simulation is that symbolic algebra systems can easily be integrated to allow formula output that can be interpreted to help find expressions for any system. While they are often simplified expressions with floating point coefficients, they can be used to investigate the behavior of a population undergoing assortative or disassortative mating in a variety of ways.

This paper will first derive the general analytical model to analyze the effects of assortative mating on two loci by showing how the difference equation for any two-locus genotype can be derived and expressed in terms of the mating covariances between multiple genotype pairs. This model will then be modified to encompass phenotype groups that contain one or more two-locus genotypes. The model can then be analyzed using a symbolic algebra system, in the case of this paper, the Symengine module in Python (Symengine-Python, v 0.11.0). Using this we can then show how metrics of the two-locus system including the single locus heterozygosity, fixation index, linkage disequilibrium, and identity disequilibrium can be derived from the two-locus genotype frequencies as they are updated each generation.

This model will then be applied to the phenotype groups from Wright’s 1921 paper to first reproduce his results for *F* and *H*_*O*_ under assortative and disassortative mating where *P*_*A*_ = *P*_*B*_ = 1*/*2, then to derive expressions for the fixation index and linkage disequilibrium under certain conditions where *P*_*A*_ = *P*_*B*_ or *P*_*A*_ = 1 − *P*_*B*_. The effects of recombination will also be analyzed in depth and the condition of *R* = 0 can be used to derive identity disequilibrium expressions for this limited case. Finally, two examples of possible assortative mating in human populations will be analyzed using the general model and available population genetic data.

## 3. A generalized assortative mating model

The key goal of population genetics is the prediction of how the genetic composition of an inter-breeding population changes over time in response to evolutionary forces and mating systems and in particular how this affects population genetic diversity. In assortative mating models with fixed allele frequencies that encompass two loci, the basic question to be answered is for any genotype *G*, what is the function *f* where *G*_*t*+1_ = *f* (*G*_*t*_) for any given two-locus genotype.

A favorable circumstance would be to have a single difference equation that could be solved analytically or graphically where *G*_*t*+1_ = *G*_*t*_ (for example see (Mickens, 2015)). From here key metrics like heterozygosity and linkage disequilibrium could be derived. However, assortative mating, while not usually affecting allele frequencies, causes complications that make this infeasible. In particular, the inbreeding like effects of reduced heterozygosity as well as the growth of linkage disequilibrium cause constant shifts in the expressions for genotype frequencies so that typical expressions used for random mating, or even consanguineous mating, cannot be simply applied to assortative mating scenarios. Therefore, the general framework has to be recursively updated for all ten genotypes until equilibrium is reached and can be analyzed.

The start of this process is identifying how to express the dynamics of assortative mating and its resulting gamete combinations to produce two-locus genotypes. In Table 2 are the expressions for all two-locus genotypes in terms of the products of gamete frequencies. These gamete frequencies have a clear expression under random mating but change in complex ways in even simple assortative mating scenarios.

**Table 2:** Two loci genotype frequencies as a function of gamete probabilities. See also (Lewontin & Kojima, 1960; Weir, 2008). Chart adopted from (Weir, 2008). Under random mating, *P*_*AB*_ = *P*_*A*_*P*_*B*_ + *D, P*_*ab*_ = (1 *− P*_*A*_)(1 *− P*_*B*_) + *D, P*_*Ab*_ = *P*_*A*_(1 *− P*_*B*_) *− D, P*_*aB*_ = (1 − *P*_*A*_)*P*_*B*_ − *D*. Note the double heterozygote (*AaBb*) genotype has two terms due to the genotype arising from either the dual gametes *AB* and *ab* or *Ab* and *aB*.

| | $AA$ | $Aa$ | $aa$ |
| --- | --- | --- | --- |
| $BB$ | $P_{AB}^2$ | $2P_{AB}P_{aB}$ | $P_{aB}^2$ |
| $Bb$ | $2P_{AB}P_{Ab}$ | $2P_{AB}P_{ab} + 2P_{Ab}P_{aB}$ | $2P_{aB}P_{ab}$ |
| $bb$ | $P_{Ab}^2$ | $2P_{Ab}P_{ab}$ | $P_{ab}^2$ |

The frequencies of the gametes can be expressed in terms of the genotypes they are derived from and then combined with the probability of mating between these genotypes to create the two-locus genotype difference equation. The probability of each gamete based on possible parental genotypes is shown in Table 3.

**Table 3:** Probabilities of gametes based on the parent genotype. The variable *R* is the recombination probability in the range 0 to 1*/*2.

| AB |  | ab |  |
| --- | --- | --- | --- |
| Parental Genotype | Probability | Parental Genotype | Probability |
| *AABB* | 1 | *aabb* | 1 |
| *AABb* | $\frac{1}{2}$ | *aaBb* | $\frac{1}{2}$ |
| *AaBB* | $\frac{1}{2}$ | *Aabb* | $\frac{1}{2}$ |
| *AB/ab* | $\frac{1}{2}(1 - R)$ | *AB/ab* | $\frac{1}{2}(1 - R)$ |
| *Ab/aB* | $\frac{1}{2}R$ | *Ab/aB* | $\frac{1}{2}R$ |

| Ab |  | aB |  |
| --- | --- | --- | --- |
| Parental Genotype | Probability | Parental Genotype | Probability |
| *AAbb* | 1 | *aaBB* | 1 |
| *AABb* | $\frac{1}{2}$ | *aaBb* | $\frac{1}{2}$ |
| *Aabb* | $\frac{1}{2}$ | *AaBB* | $\frac{1}{2}$ |
| *AB/ab* | $\frac{1}{2}R$ | *AB/ab* | $\frac{1}{2}R$ |
| *Ab/aB* | $\frac{1}{2}(1 - R)$ | *Ab/aB* | $\frac{1}{2}(1 - R)$ |

The key to creating the difference equation for genotype probabilities is realizing that the frequency of every two-locus genotype can be represented as the product of the frequencies of two gametes and that these products of gamete frequencies themselves can be represented as the weighted sum of the probabilities of matings between multiple genotype pair combinations.

For example, for the genotype *AABB*, its frequency is *P*_*AABB*_ = *P*_*AB*_*P*_*AB*_. The total frequency for the gamete *AB* is *P*_*AB*_ = *P*_*AABB*_ + 1*/*2(*P*_*AABb*_ + *P*_*AaBB*_) + 1*/*2(1 *− R*)*P*_*AB/ab*_ + 1*/*2*RP*_*Ab/aB*_. The frequency *P*_*AABB*_ thus can also be expressed as (*P*_*AABB*_ + 1*/*2(*P*_*AABb*_ + *P*_*AaBB*_) + 1*/*2(1 *− R*)*P*_*AB/ab*_ + 1*/*2*RP*_*Ab/aB*_)*×*(*P*_*AABB*_+1*/*2(*P*_*AABb*_+*P*_*AaBB*_)+1*/*2(1 *− R*)*P*_*AB/ab*_+1*/*2*RP*_*Ab/aB*_) where the product not only multiplies the term coefficients but also indicates matings between crossed genotypes.

The first term would be *P*_*AABB×AABB*_, where *×* represents a genotype mating. The next terms would be 1*/*2*P*_*AABB×AABb*_, 1*/*2*P*_*AABB×AaBB*_, 1*/*2(1*− R*)*P*_*AABB×AB/ab*_ etc. The final expression for *P*_*AABB*_, multiplied over all terms, is given by

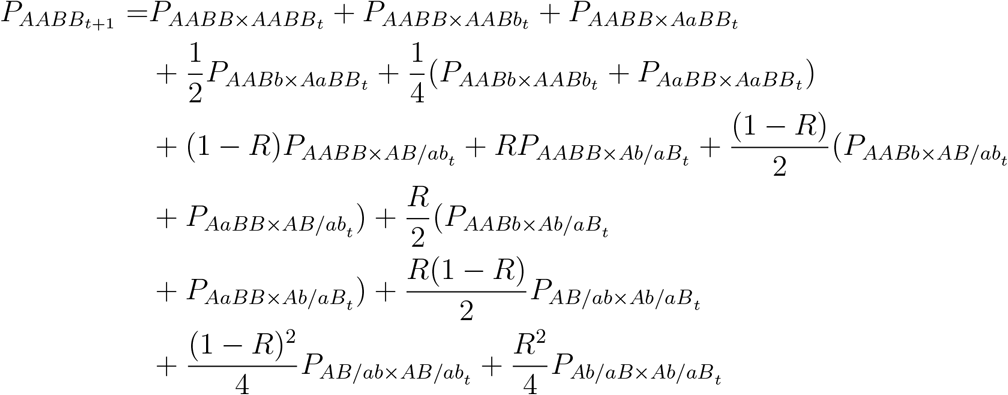

Likewise, given *P*_*ab*_ = *P*_*aabb*_ + 1*/*2(*P*_*Aabb*_ + *P*_*aaBb*_) + 1*/*2(1 *− R*)*P*_*AB/ab*_ + 1*/*2*RP*_*Ab/aB*_, the double heterozygote *AB/ab* frequency *P*_*AB/ab*_ = 2*P*_*AB*_*P*_*ab*_ can be shown by

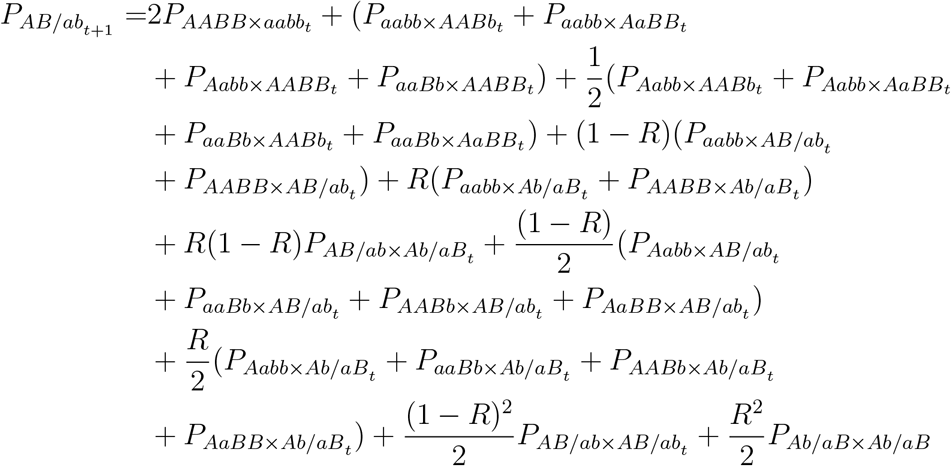

Similar expressions for the other eight two-locus genotypes can be easily derived using the methods above and Tables 2 and 3. To analyze these expressions further, it is necessary to derive an expression for the probability of two genotypes mating. For two genotypes, *G*_1_ and *G*_2_, their mating probability 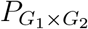 can be expressed in a manner similar to linkage disequilibrium. The covariance of mating between the two genotypes is

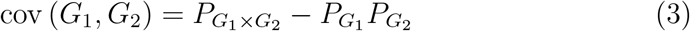

So the probability of mating is given by

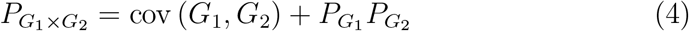

All mating probabilities in the genotype difference equations can be substituted with similar expressions. However, first the overall resulting expression can be simplified. When there is random mating all mating covariances are equal to zero thus 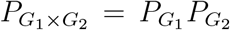. Assuming we are starting from the first generation of assortative mating where *t* = 0 and the two-locus genotypes can be derived from the expressions in Table 2 using the gamete expressions shown in the table legend, what we find, using the symbolic computation of Symengine, for each genotype is that when all covariances are zero, the resulting sum over all genotypes 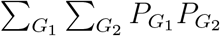 gives the two-locus genotype frequency as expected when only recombination is present as a factor in changing genotype frequencies. To use *AABB* as an example, the resulting expression becomes

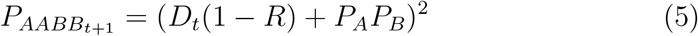

Of course when *R* = 0 or *D* = 0, the genotype frequency is at a steady state and does not change. Therefore, all of the genotype difference equations can be expanded to be the sum of the random mating case plus the effect of all covariances of mating. Restating the expression for *AABB*

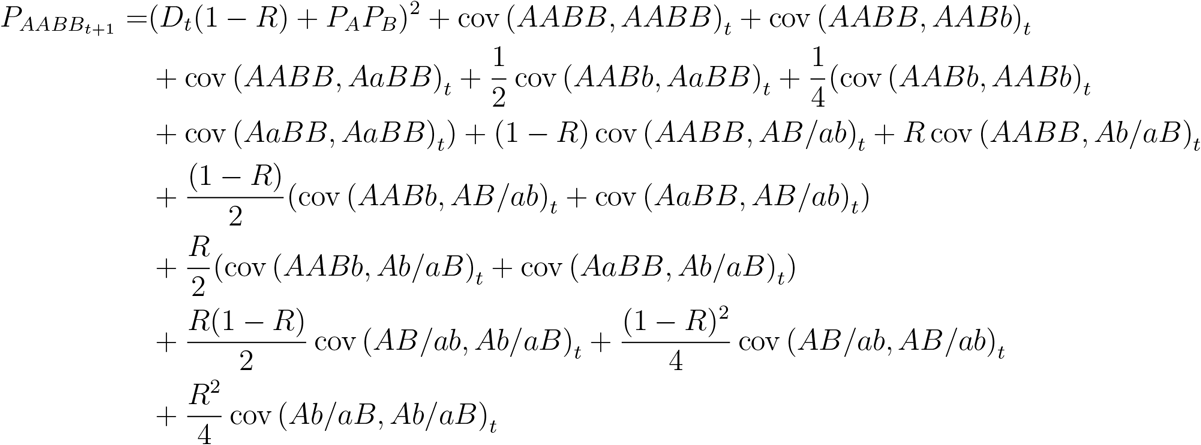

The mating covariances between genotypes can be derived from theoretical or natural observed mating correlations, however, as the genotype frequencies change over time, the covariances would have to be adjusted to compensate for the new genotype variances. Another solution is to use a normalized covariance, similar to Lewontin’s *D*^*′*^ in linkage disequilibrium, to have a constant factor that can be used to measure the relative strength of assortative mating between genotypes. For this purpose we will use Hedrick’s *A*, first defined in (Hedrick, 2017). While *A* in that paper was defined as the proportion of matings that were within phenotype groups in a population, we will show a definition of *A* below behaves similarly to the example Hedrick used. Here *A* is defined for assortative mating between genotypes as

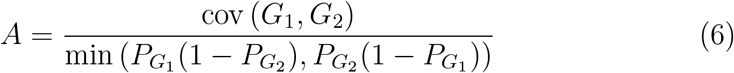

and for disassortative matings between genotypes as

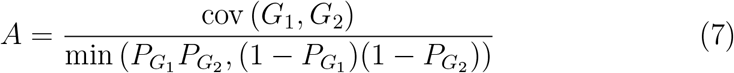

For phenotypes that preferentially mate with each other, *A >* 0, while *A <* 0 for phenotypes that disproportionately avoid mating. While it will not be shown in the paper, the covariances cov (*G*_1_, *G*_2_) in the genotype difference equations can be replaced with the appropriate expressions containing *A* and the maximum covariance. After establishing these definitions, one can use arbitrary allele frequencies, starting linkage disequilibrium values, and values of assortative/disassortative mating to recursively simulate the results in a population undergoing assortative mating to find equilibrium states or even intermediate values. Given these recursions are deterministic, they assume an infinitely sized population and do not require statistical adjustments or bounds in their analysis. This means they will always give identical results under the same conditions. However, this means they can give inaccurate results for small populations where factors such as drift or consanguineous mating may come into play.

## 4. Incorporating phenotype groups into the model

The model shown in the previous section assumes that each genotype can be treated as a separate phenotype. However, this is often not the case, particularly in the Wright model which will be analyzed in this paper. Fortunately, it is easy to accommodate phenotype groups into the analysis. Given phenotype groups *X*_1_ and *X*_2_ which each have genotypes *G*_1_ and *G*_2_ as (non-exclusive) members and assuming not only that the mating covariances are between phenotypes but when phenotypes mate, the underlying genotypes are selected at random from those in the phenotype group, the mating probability between genotypes can be restated as below.

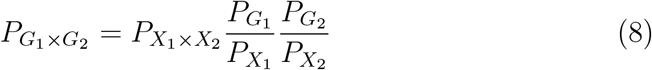

The phenotype frequencies are the sum of the frequencies of all genotypes that are expressed as that phenotype. Furthermore, the probability of phenotype group mating can be defined similar to equation 4.

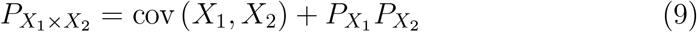

The normalized assortative mating coefficient *A* can be redefined similarly for assortative and disassortative mating between phenotype groups as

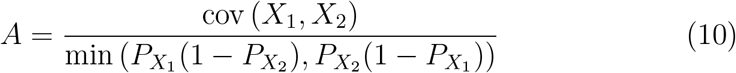

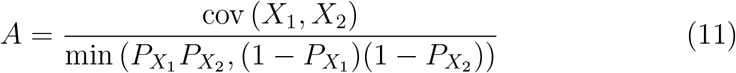

## 5. Accommodating allele frequency changes

While standard assortative mating models generally do not result in allele frequency changes, the same is not true for disassortative mating where it has been argued that disassortative mating often inevitably leads to allele frequency changes in most cases. This phenomenon and links to selective mating were discussed in (Spencer, 1992). However, since the only disassortative models investigated are typically the simple one where *P*_*A*_ = *P*_*B*_ = 1*/*2, these conditions are not widely discussed.

As part of the investigation of disassortative mating, the model was adapted to incorporate allele frequency changes. In short, the value of 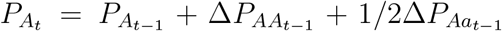. This can be expanded using the two-locus genotypes as

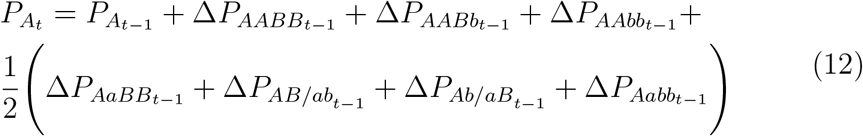

Analysis of the allele frequency changes for various conditions of disassortative mating show only a narrow set of conditions where allele frequencies do not change when disassortative mating is present.

## 6 Measured genetic aspects of assortative mating populations

Using the previously derived expressions we can now subject the Wright model to analysis through recursive calculations of the difference equations for each two-locus genotype. Using the phenotype groups shown in Table 1 and assuming a the alleles are at linkage equilibrium at *t* = 0, calculations of the frequencies of two-locus genotypes are carried out at each time step and updated. Initial allele frequencies remain constant under assortative mating but change in many instances of disassortative mating.

There are several key metrics amongst one or two-locus genotypes that change under assortative mating. Not only single locus heterozygosity, but linkage disequilibrium, Cockerham and Weir’s composite linkage disequilibrium, and identity disequilibrium.

### 6.1 Single locus heterozygosity and the fixation index

Single locus observed heterozygosity *H*_*O*_ = *P*_*Aa*_ is calculated by summing the corresponding two-locus genotypes

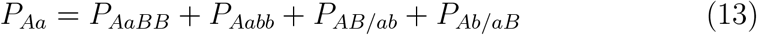

The corresponding fixation index is calculated as

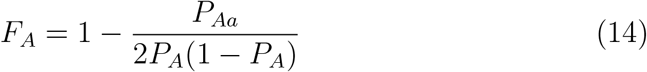

### 6.2 Linkage disequilibrium

There are two well-known expressions to calculate linkage disequilibrium. First is using gamete frequencies such as *D* = *P*_*AB*_–*P*_*A*_*P*_*B*_. Second is using the frequencies of the double heterozygotes *D* = *P*_*AB/ab*_–*P*_*Ab/aB*_. The first will be used in our analysis. The second is not valid when reduced heterozygosity due to inbreeding or other non-random mating occurs. It is replaced by composite linkage disequilibrium (Cockerham & Weir, 1973) which will also be calculated separately.

The gamete frequency (after recombination) *P*_*AB*_ is given by

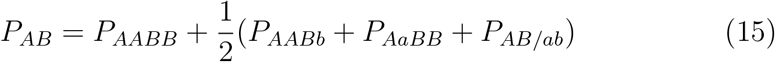

The maximum values of *D* given allele frequencies, *D*_max_ are given below. As standard, Lewontin’s *D*^*′*^ is given by *D*^*′*^ = *D/D*_max_.

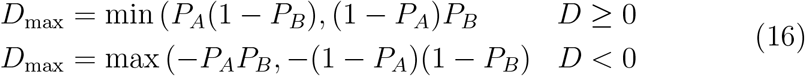

Finally, the correlation coefficient between alleles is given by

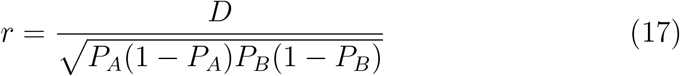

#### 6.2.1 Composite linkage disequilibrium

In (Cockerham & Weir, 1973), it was noted that the difference between double heterozygotes *P*_*AB/ab*_*− P*_*Ab/aB*_ does not yield just the value of linkage disequilibrium *D* in systems of inbreeding as in the random mating scenario. Rather, it represents a composite linkage disequilibrium that takes into account the structure of the inbreeding in the population using two of what they termed “descent coefficients”. Only when inbreeding was absent did the value revert back to *D*. Composite linkage disequilibrium will be analyzed by looking at this difference. It is found, however, that often its value is zero under the Wright model since under most scenarios the double heterozygotes have a nearly equal frequency at equilibrium (or both are absent).

### 6.3 Identity disequilibrium

Identity disequilibrium is a property of two-locus systems with inbreeding that increases the frequencies of double homozygotes and double heterozygotes, while decreasing the frequency of homozygote-heterozygotes. In consanguineous mating, it is based on both the amount of linkage and the nature of parental consanguinity. It is also the correlation between heterozygosity or between homozygosity at two loci, with heterozygosity or homozygosity co-existing at both loci more often than random co-occurrence would suggest.

Often designated as *η*_11_ its effect on the two-locus genotype frequencies, when only inbreeding and not linkage disequilibrium is present, is shown in Table 4. See also (Haldane, 1949; Cockerham & Weir, 1973; Smith, 2023) for details.

**Table 4:**
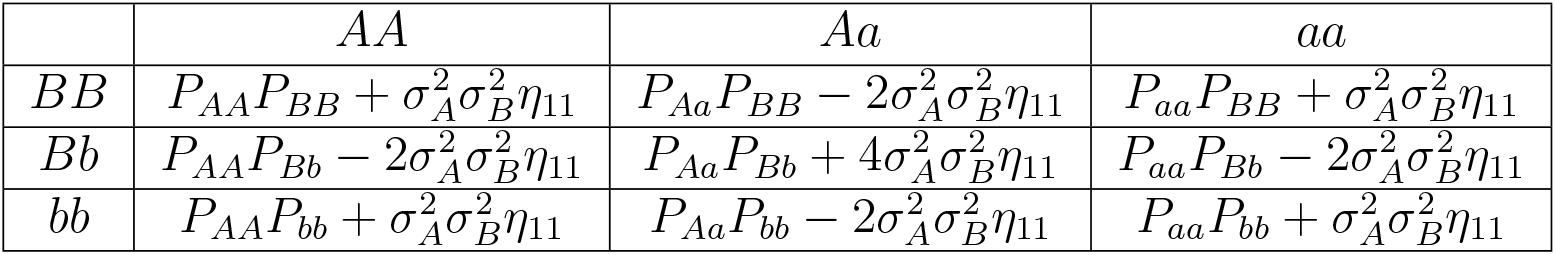
Two loci genotype frequencies under inbreeding at linkage equilibrium taking identity disequilibrium into account. For original explanations and derivations see (Haldane, 1949), though he uses *ϕ* for *η*_11_. The variance of each locus is represented by 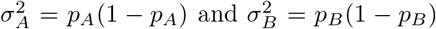. The variable *η*_11_ is the identity disequilibrium. Per the regular results of inbreeding, 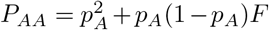, *P*_*Aa*_ = 2*p*_*A*_(1 *−p*_*A*_)(1 *−F*), *P*_*aa*_ = (1 *− p*_*A*_)^2^ + *p*_*A*_(1 *− p*_*A*_)*F*, 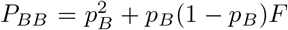, *P*_*Bb*_ = 2*p*_*B*_(1 *− p*_*B*_)(1 *− F*), *P*_*bb*_ = (1 *− p*_*B*_)^2^ + *p*_*B*_(1 *− p*_*B*_)*F*. If the loci are unlinked *η*_11_ = 0 and if they are completely linked *η*_11_ = *F* (1 *− F*). Note double heterozygotes have a factor of four for identity disequilibrium due to the two types of combining gametes (*AB* and *ab* or *Ab* and *aB*) that create double heterozygotes and the possibility of each gamete being on the paternal or maternal contributed chromosome. Similarly homozygotes-heterozygotes have a factor of two in relation to identity disequilibrium since each genotype can appear two different ways depending on its contribution from maternal or paternal chromosomes.

| | $AA$ | $Aa$ | $aa$ |
| --- | --- | --- | --- |
| $BB$ | $P_{AA}P_{BB} + \sigma_A^2\sigma_B^2\eta_{11}$ | $P_{Aa}P_{BB} - 2\sigma_A^2\sigma_B^2\eta_{11}$ | $P_{aa}P_{BB} + \sigma_A^2\sigma_B^2\eta_{11}$ |
| $Bb$ | $P_{AA}P_{Bb} - 2\sigma_A^2\sigma_B^2\eta_{11}$ | $P_{Aa}P_{Bb} + 4\sigma_A^2\sigma_B^2\eta_{11}$ | $P_{aa}P_{Bb} - 2\sigma_A^2\sigma_B^2\eta_{11}$ |
| $bb$ | $P_{AA}P_{bb} + \sigma_A^2\sigma_B^2\eta_{11}$ | $P_{Aa}P_{bb} - 2\sigma_A^2\sigma_B^2\eta_{11}$ | $P_{aa}P_{bb} + \sigma_A^2\sigma_B^2\eta_{11}$ |

In the calculations, the identity disequilibrium is calculated by analyzing the correlation between *AA* and *BB* homozygotes at the two loci using the following expression (Smith, 2023).

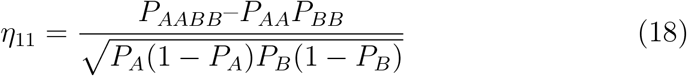

## 7. Key model results

Using the phenotype groups of (Wright, 1921), the model will be used to analyze the equilibrium values of population genetic variables. First, in order to verify the validity of the results, the model was analyzed using the traditional allele frequencies of *P*_*A*_ = *P*_*B*_ = 1*/*2.

### 7.1. Assortative Mating for P_*A*_ = *P*_*B*_ = 1*/*2

This is the model most applicable to almost all past work on assortative mating. Assortative mating was established where matings occurred between individuals of the same phenotype based on a given *A* value assumed for all groups in the population. Matings between phenotypes had a propensity of*− A*. The equations were recursively solved until the solutions approximated equilibrium (30 generations). The loci were considered unlinked so *R* = 1*/*2. The equilibrium heterozygosity, to be termed *H*_*∞*_, ended up being expressed by the program in terms of *H*_*∞*_ = 1*/*2–*CA*. Here *C* is a constant that changed based on the value of *A*. However, in all cases it was found that when *H*_*E*_ = 1*/*2 is factored out, *C/H*_*E*_ = 1*/*(4 *−* 3*A*). In other words, this fits the original expression by Wright and also shows that in the case of *P*_*A*_ = *P*_*B*_ = 1*/*2, *A* = *m*. Thus both the observed heterozygosity and the value of the fixation index, *F*, fit the original solution of Wright.

Next was to investigate the values of linkage disequilibrium. Similar to heterozygosity, it was given by a value of *D*_*∞*_ = *C*_0_–*C*_1_*A*. Where *C*_0_ and *C*_1_ are constants. At *A* = 1, *C*_0_ = 1*/*2 and *C*_1_ = 0.25. However, unlike heterozygosity where the first term in the difference always equaled the expected heterozygosity, *C*_0_ varied as well as *C*_1_ with *A*. However, plotting *D*_*∞*_ versus *H*_*∞*_ revealed a linear and consistent relationship where *D*_*∞*_ = –*H*_*∞*_*/*2. Given *H*_*∞*_ = 1*/*2(1 *−F*) and *F* = *A/*(4 *−* 3*A*), the expression for linkage disequilibrium is thus

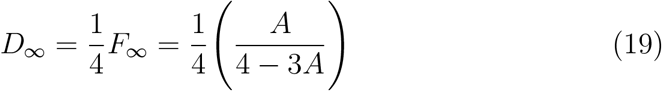

One interesting aspect of the linkage disequilibrium change when viewed in successive generations, is that under all values of *A*, there is no linkage disequilibrium after the first generation of assortative mating. This was hinted at by Wright when he described the correlation between gametes in the first generation as zero. Only the inbreeding like effect of reduced heterozygosity is present. In the first generation, the value of *F* is dependent on *A* so that

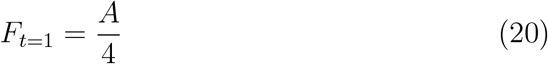

In the second generation of mating, heterozygosity continues to decline but now linkage disequilibrium increases as well.

### 7.2. Disassortative Mating for P_*A*_ = *P*_*B*_ = 1*/*2

Similar to the above, disassortative mating was tested using Wright’s model and the disassortative matings outlined in Table 1. Here the within phenotype mating propensity becomes *−A*. One of the first observations was equilibrium values are reached much more rapidly for disassortative mating—in 10 generations or less. The symbolic output of the heterozygosity was similar with an additive term instead of a subtracting one indicating the increase in heterozygosity. In (Wright, 1921), it was noticed that disassortative mating could be handled by making the correlation negative and we find that again *A* = *m* in this expression. The equilibrium fixation index for disassortative mating, which is negative, is

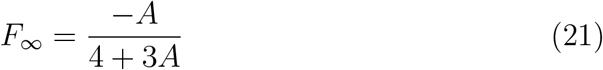

Final heterozygosity, which increases, is given by

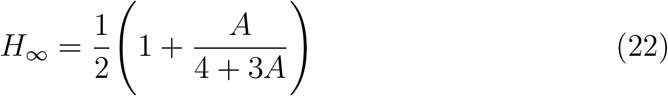

Linkage disequilibrium is negative in disassortative mating and its symbolic relationship was similar to that in assortative mating. Again plotting values against heterozygosity one finds the same relationship.

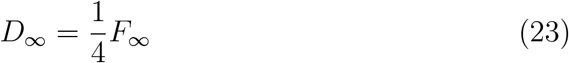

Note that the smallest value for *F*_*∞*_ for disassortative mating in this context where *A* = 1 is *F*_*∞*_ = *−*1*/*7. Therefore the minimum value for linkage disequilibrium is *D*_*∞*_ = *−*1*/*28 *≈ −*0.0357.

### 7.3. Assortative Mating for P_*A*_ = *P*_*B*_

After establishing the results for the case of both loci having a frequency of 1*/*2, we now look at the case where *P*_*A*_ = *P*_*B*_ for any arbitrary value of *P*_*A*_. Following the methods of looking at the expression output for *F* and *D* that were used for the previous case, we find that the results for *P*_*A*_ = *P*_*B*_ = 1*/*2 are generalized for all assortative mating where *P*_*A*_ = *P*_*B*_. First, the equilibrium fixation index *F* is still

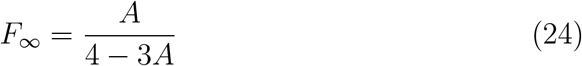

Thus this expression can be used directly to find the equilibrium heterozygosity,

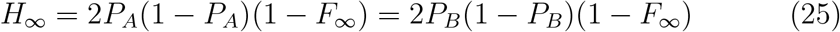

Similar to the case of 1*/*2, the linkage disequilibrium is zero in the first generation. The value of the fixation index after the first generation remains the same as equation 20. In addition, the ratio between the value of the fixation index in the first generation and the equilibrium fixation index is the same for all *P*_*A*_ = *P*_*B*_.

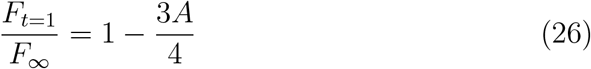

Linkage disequilibrium finds a similar generalization as well, again in terms of *P*_*A*_

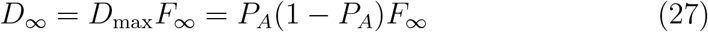

This interesting result solidifies what was only hinted at when *P*_*A*_ = *P*_*B*_ = 1*/*2, namely that linkage disequilibrium from assortative mating when *P*_*A*_ = *P*_*B*_ is directly proportional to the inbreeding depression in the population which is 2*P*_*A*_(1 *− P*_*A*_)*F*_*∞*_*d*_1_. The value *d*_1_ is the dominance effect size that is equal to the difference between the mean for a phenotypic trait between the two homozygotes. Also since *D*_max_ = *P*_*A*_(1 *− P*_*A*_) and *r* = *D/*(*P*_*A*_(1 *− P*_*A*_)) when *P*_*A*_ = *P*_*B*_, *D*^*′*^ = *r* and we also have the relationship

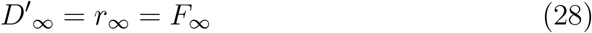

Therefore, the equilibrium linkage disequilibrium correlation is always equal to the equilibrium fixation index when *P*_*A*_ = *P*_*B*_.

One interesting aspect of the expression *D*^*′*^ = *r* is that it signifies the linkage disequilibrium generated by assortative mating under these conditions can always have a range of [0, 1] for *r* (VanLiere & Rosenberg, 2008). Likewise, the quantity 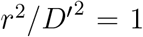 indicates that the linkage disequilibrium is completely linear, a consequence of allele frequency matching at both loci, and all dependence can be explained by *r* (Smith, 2020). This is a condition that we will see is only met in one other context: for disassortative mating when *P*_*A*_ = 1 *−P*_*B*_. These two contexts are also the only ones where this research found widely applicable closed form expressions for linkage disequilibrium.

### 7.4. Assortative Mating for *P*_*A*_ = 1 *− P*_*B*_

The case of *P*_*A*_ = 1 *− P*_*B*_ is the only other case where assortative mating results in the same heterozygosity frequency and fixation index at both loci. Therefore, it is the next easiest to model and determine. From the outset it is clear that the Wright equation for *F*_*∞*_ that works for *P*_*A*_ = *P*_*B*_ does not fit the data, nor does the expression for *D*.

There is no closed form derivation of *F*_*∞*_ in this paper but we are able to use the special cases of *A* = 0 and *A* = 1 to find a close approximation for *F*_*∞*_. The formulaic output for *H*_*∞*_ for all values of *A* is similar as before and is given by (at locus A) *H*_*∞*_ = 2*P*_*A*_(1 *− P*_*A*_)–*C*_2_*A*. A value is given for *C*_2_ by the symbolic computation even when *A* = 0 allowing us to derive an expression for *F*_*∞*_. Assuming the general form of *F*_*∞*_ is similar to the definition of *F*_*∞*_ where *P*_*A*_ = *P*_*B*_, when *A* = 0,

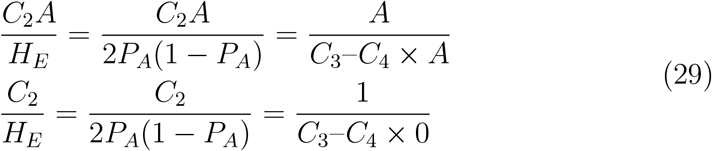

In the expression above, the second equation is the form of *F*_*∞*_ for *P*_*A*_ = *P*_*B*_ where *C*_3_ = 4 and *C*_3_ = 3.

This allows us to define *C*_3_ as

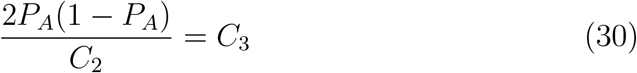

A plot of *C*_3_ for values of *P*_*A*_ is given in Figure 2 shows a roughly quadratic function ranging from approximately 2 where the minor allele frequency at each locus is small (0.01) to 4, as expected, where *P*_*A*_ = *P*_*B*_ = 1*/*2. In fact, the maximum and minimum values, as well as many intermediate values, approximate 2*N*_*e*_ where *N*_*e*_ is the effective number of alleles at each locus (Kimura & Crow, 1964).

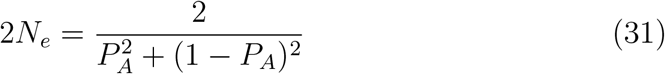

As shown in Figure 2, Equation 31 closely approximates the values. Next, in order to solve for *C*_4_, we will utilize the situation where *A* = 1. Using Equation 31 for *C*_3_ we can solve *C*_4_ using

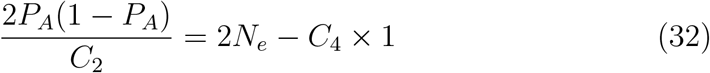

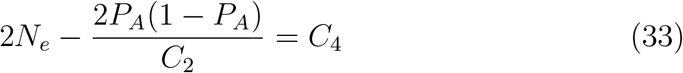

Figure 3 shows a similar relationship to that for *C*_3_ except the approximation here is 2*N*_*e*_ *−* 1. Combining this presents a reasonable approximation for *F*_*∞*_ where *P*_*A*_ = 1 *− P*_*B*_.

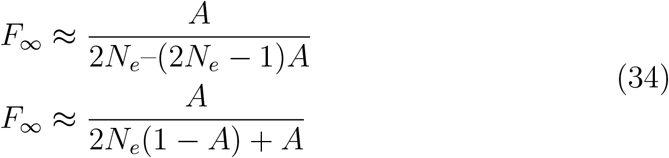

Measured values of *F*_*∞*_ and those calculated with Equation 34 are shown in Figure 4.

**Figure 1.**
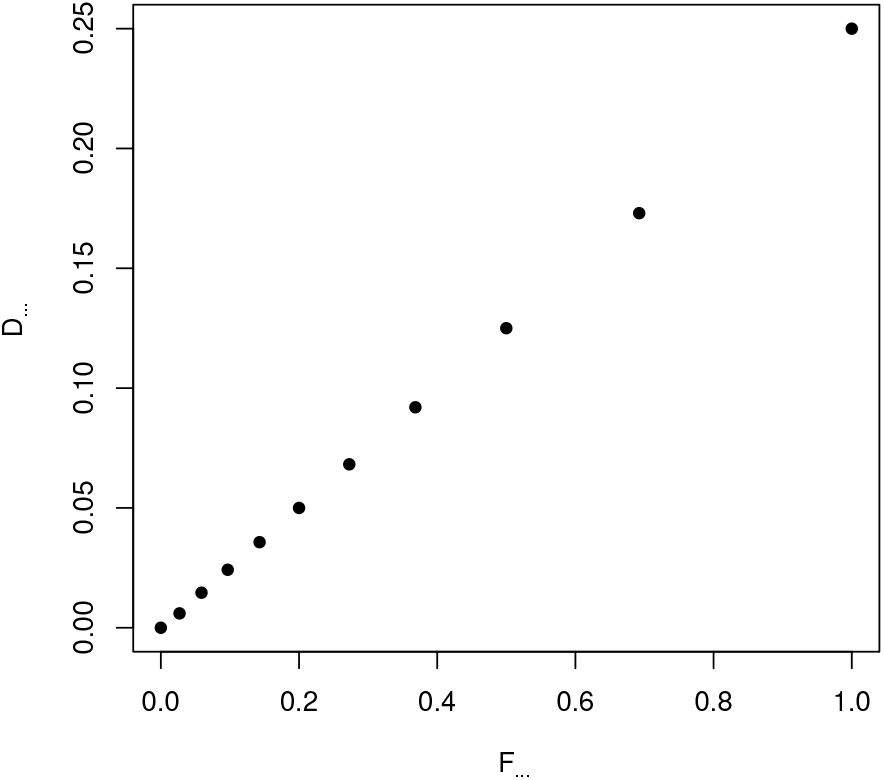
A plot of the values of *D*_*∞*_ vs. *F*_*∞*_ for *P*_*A*_ = *P*_*B*_ = 1*/*2. The fitted slope is 1/4.

**Figure 2.**
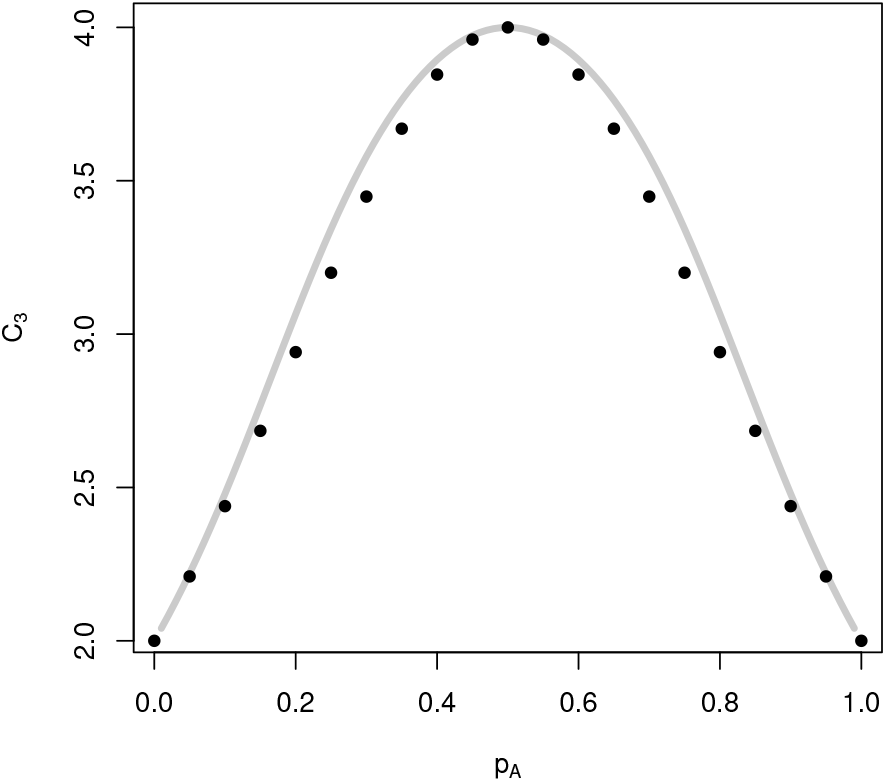
A plot of the values for the imputed coefficient *C*_3_ versus *P*_*A*_. Simulation values are the gray line, calculated values of 2*N*_*e*_ are the black dots.

**Figure 3.**
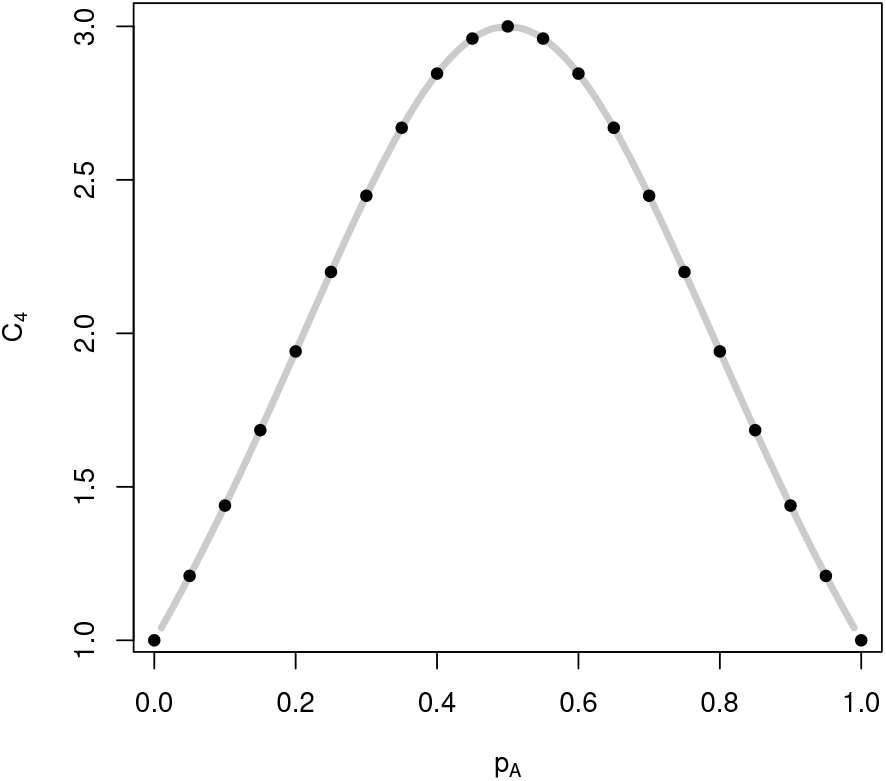
A plot of the values for the imputed coefficient *C*_4_ versus *P*_*A*_. Simulation values are the gray line, calculated values of 2*N*_*e*_ *−* 1 are the black dots.

**Figure 4.**
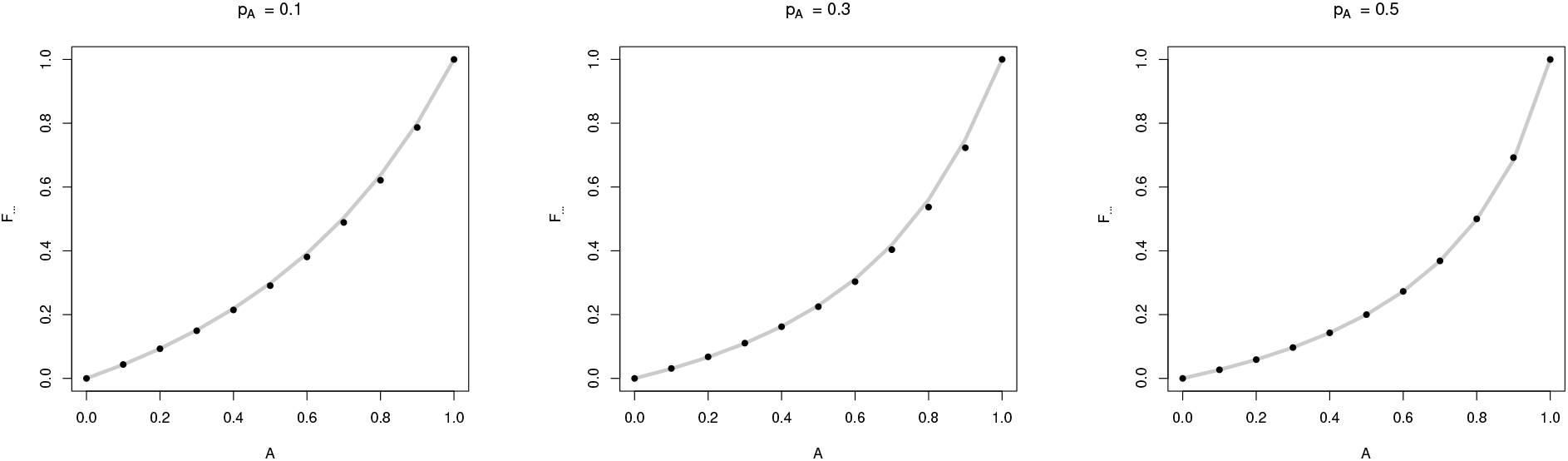
Plots for the value of *F*_*∞*_ when *P*_*A*_ = 1 *− P*_*B*_ at both loci using the actual (solid gray line) simulation values and values estimated using Equation 34 (black dots). Values of *P*_*A*_ shown are *P*_*A*_ = 0.1, *P*_*A*_ = 0.3, and *P*_*A*_ = 0.5.

Linkage disequilibrium, however, does not find such a clear resolution. Unlike where *P*_*A*_ = *P*_*B*_, *D*^*′*^*≠ r* and the range of *r* is constricted and even low for most assortative matings for *P*_*A*_ = 1 *− P*_*B*_ *≠* 1*/*2. In fact, when one of the allele frequencies is low, such as 0.3 or lower, *r* only reaches small values and most dependence is with *D*^*′*^ indicating that the linkage disequilibrium is generally nonlinear since the allele frequencies do not match (Smith, 2020).

One consistent pattern, however, is that *D*_*∞*_ *> D*_max_*F*_*∞*_ which indicates more linkage disequilibrium than expected by an equivalent expression for the *P*_*A*_ = *P*_*B*_ case. No clear and general expression was found using symbolic algebra or equation fitting approximations. Therefore, surprisingly the clearest expressions for *D* for assortative mating remain those when *P*_*A*_ = *P*_*B*_.

### 7.5. Disassortative Mating for P_*A*_ = *P*_*B*_*≠*1*/*2

While the case of assortative mating *P*_*A*_ = *P*_*B*_ is a straightforward extrapolation from the *P*_*A*_ = *P*_*B*_ = 1*/*2 frequency case, disassortative mating is much more complex. In short, for all combinations of *P*_*A*_ = *P*_*B*_ ≠1*/*2, there is a change in the allele frequencies of the population as disassortative mating occurs. There is no readily clear or consistent pattern evident and allele frequency changes range from relatively minor to fixation of an allele at a locus. With such changes the standard expressions for straightforward analysis of the system become those of selective mating more than those of assortative mating (often a subset as shown in (Spencer, 1992)). While this is definitely an avenue for investigation it will not be covered in the paper due to the limits of the applied methodology.

### 7.6. Disassortative Mating for P_*A*_ = 1 *− P*_*B*_

Unlike the case of *P*_*A*_ = *P*_*B*_ ≠ 1*/*2, disassortative mating where *P*_*A*_ = 1 *− P*_*B*_ is the only condition where disassortative mating does not change allele frequencies. Simulation amongst various allele frequency values for each locus across the entire range from 0.01 to 0.99 consistently show that only when *P*_*A*_ = 1 *− P*_*B*_ do allele frequencies stay fixed for all generations of disassortative mating. In Figure 5, a contour plot shows the allele frequency changes Δ*P*_*A*_ (equilibrium allele frequency after disassortative mating minus starting allele frequency) caused by disassortative mating for several values of *A*. The line of *P*_*A*_ = 1 *− P*_*B*_ clearly shows no change in allele frequency while other regions show various values of increases or decreases in allele frequency caused by disassortative mating.

**Figure 5.**
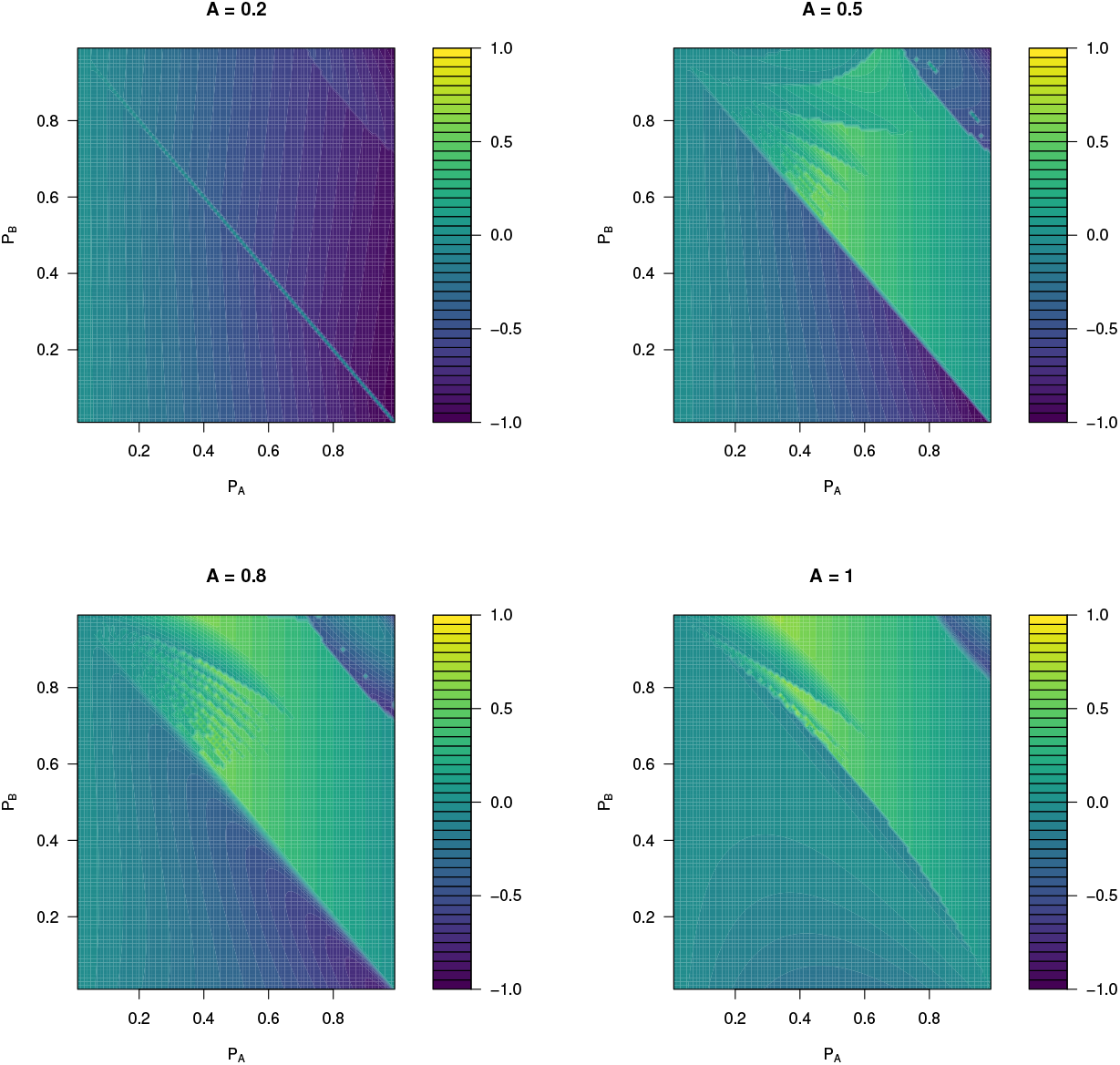
Contour plots of Δ*P*_*A*_ due to disassortative mating depending on the initial values (*t* = 0) of *P*_*A*_ (x) and *P*_*B*_ (y) and *A*.

However, the relationship describing *F*_*∞*_ is not as simple as making *A* negative that is the case where *P*_*A*_ = *P*_*B*_ = 1*/*2. Rather the relationship is again approximated using the cases of *A* = 0 and *A* = 1, similar to the previous example for assortative mating when *P*_*A*_ = 1 *− P*_*B*_, to yield the following approximation.

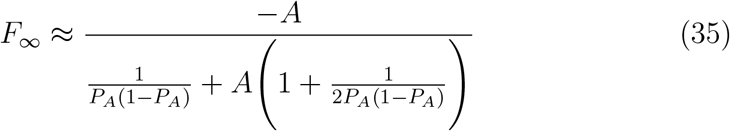

Fits to data are shown in Figure 6.

**Figure 6.**
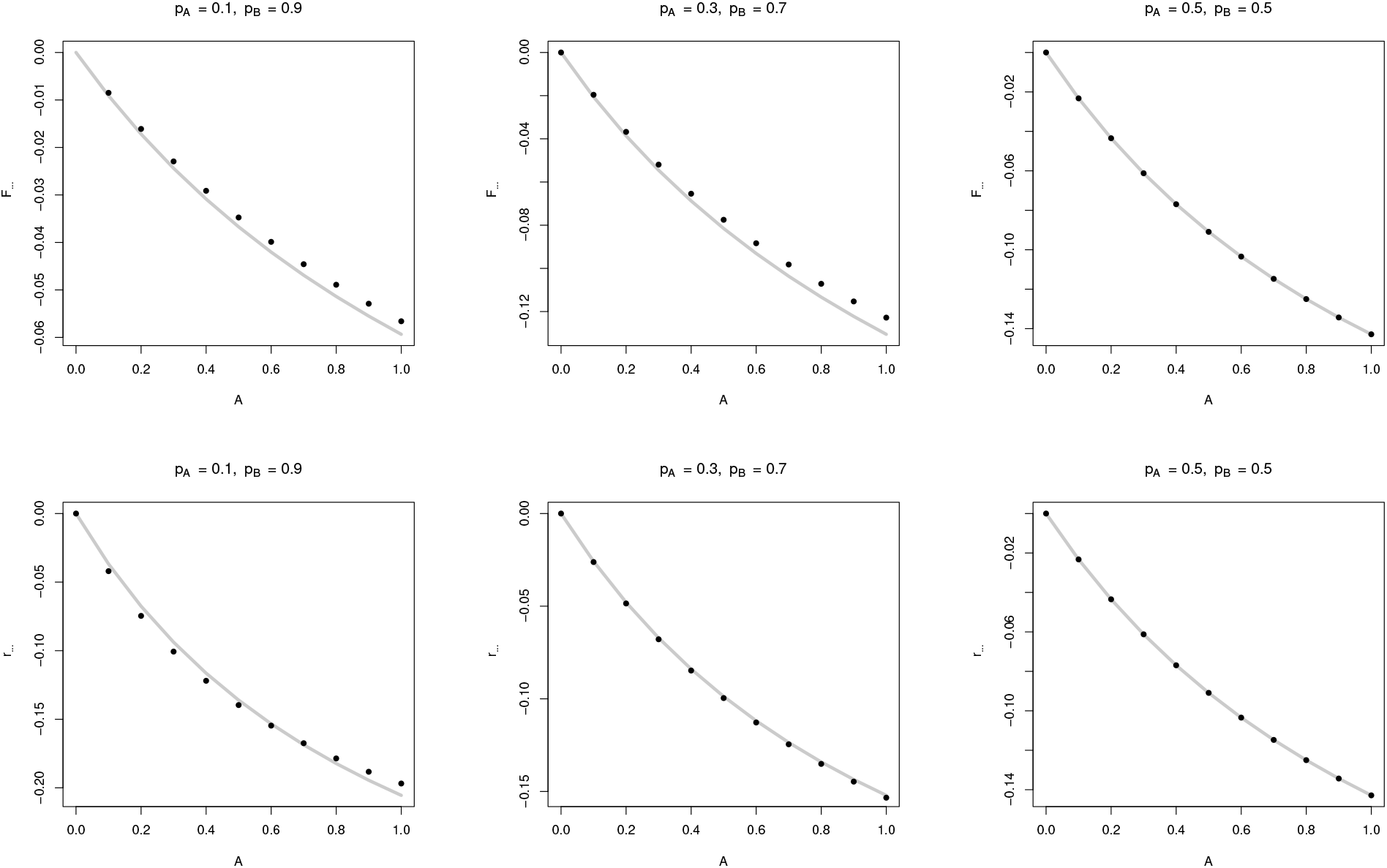
Comparisons of data to approximate expressions for *F*_*∞*_ (top row) and *r*_*∞*_ (bottom row) for disassortative mating when *P*_*A*_ = 1 *− P*_*B*_. Actual simulation data uses solid gray lines and calculated values are shown as black dots. Values of *F*_*∞*_ are estimated with Equation 35 and the values of *r*_*∞*_ are estimated with Equation 36. Values of *P*_*A*_ shown are *P*_*A*_ = 0.1, *P*_*A*_ = 0.3, and *P*_*A*_ = 0.5.

As stated earlier, negative linkage disequilibrium meets the full conditions for linearity, 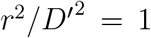 and thus a full range of *r* from [-1,0] is possible though *r* does not approach -1 in any combination of allele frequencies and *A*. Linkage disequilibrium has a fitted expression, discovered with the help of the Nutonian Eureqa equation fitting software (Eureqa, v1.12.1), that is similar to the expression for *F*_*∞*_ where *P*_*A*_ = *P*_*B*_ = 1*/*2 but it does not have a clear relationship to the value of *F*_*∞*_. Given *r*_*∞*_ *<* 0 the value of 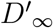 is 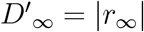.

The approximate expressions for *r*_*∞*_ and *D*_*∞*_ are

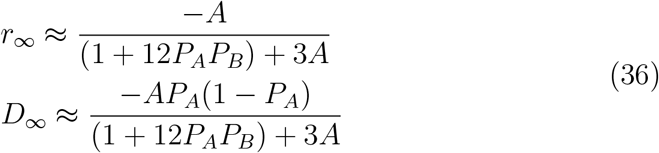

The value of the denominator in the expression depends on the variance at the loci under consideration and converges to Equation 21 when *P*_*A*_ = *P*_*B*_ = 1*/*2. Fits to data are shown in Figure 6 as well.

## 8. Assortative Mating for *P*_*A*_ *≠ P*_*B*_ *≠*1 *− P*_*B*_

Outside of the cases above where the variance at both loci are the same, the results of assortative mating become more complex still. The equilibrium values of *F*_*∞*_ are different at each locus even though they may be similar in magnitude. They are only identical when *A* = 0 and *F*_*∞*_ = 0 and *D*_*∞*_ = 0 or *A* = 1 and *F*_*∞*_ = 1 and *D*_*∞*_ = *D*_max_ as (Ghai, 1973) intimated for *A* = 1 using two-locus genotype frequencies.

The difference in *F*_*∞*_ at each locus is proportional to the difference in the size of the variance at each locus. Therefore the difference in *F*_*∞*_ increases as the difference in the variance at each locus increases. Likewise, while 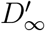 ranges from 0 to 1 as *A* ranges from 0 to 1 for any combination, *r* has a relatively small value except when the loci variances are close or equal. Beyond these facts, a consolidated and general group of expressions for assortative mating has yet to be found.

## 9. The effects of recombination on the approach to equilibrium

In early work on assortative mating, Robbins (Robbins, 1918) noted that complete linkage between loci yielded results consistent with linkage disequilibrium being zero. In (Hedrick, 2017), it was noted that in simulations the final value of linkage disequilibrium was not affected by the linkage between loci. However, the rate of approach to this equilibrium increased as the linkage between loci decreased. The same results are found here, with some modification, and will be shown for assortative and disassortative mating.

### 9.1. Effects of recombination under assortative mating where P_A_ = *P*_*B*_

The increase in linkage disequilibrium across time can be shown by curves for differing degrees of linkage between loci. In Figure 7 the curves for *P*_*A*_ = *P*_*B*_ = 1*/*2 are shown for *A* = 1 and values of *R* ranging from 0 to 1*/*2. One of the first findings is confirmation that linkage disequilibrium is not formed by assortative mating when *R* = 0. This is intuitive since the frequency changes in haplotypes cannot occur without recombination when allele frequencies are fixed. Reduction of heterozygosity at each locus does still occur when *R* = 0 but it is not as extensive and heterozygotes and double heterozygotes do not completely disappear, even if *A* = 1.

**Figure 7.**
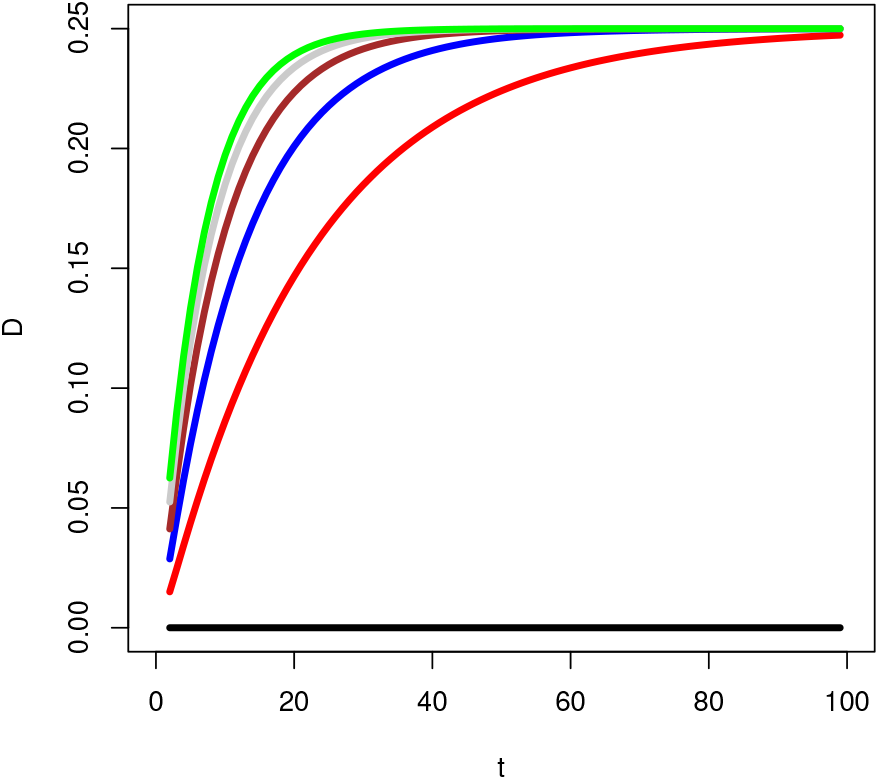
Plot of the increase in linkage disequilibrium by generation (*t >* 1) for various values of recombination between the loci where *P*_*A*_ = *P*_*B*_ = 1*/*2 and *A* = 1. The values of *R* are 0 (black), 0.1 (red), 0.2 (blue), 0.3 (brown), 0.4 (gray), and 0.5 (green).

The implication is that linkage disequilibrium between linked loci can be nearly ruled out as having arisen from assortative mating. Linkage disequilibrium will not be created between linked loci in cases where the allele frequencies are not changing. However, the decline of heterozygosity still continues, though its ultimate value is higher than if recombination was present. In the simplest case, where *A* = 1, *F*_*∞*_ = 1*/*2 and the final heterozygosity is one half of its value at Hardy-Weinberg equilibrium.

A clear way to compare the curves under different values of linkage is to use the *R* = 1*/*2 case as a baseline and compare other values of linkage to it. For all *P*_*A*_ = *P*_*B*_, the rate of increase of linkage disequilibrium over time for unlinked loci has an approximate expression.

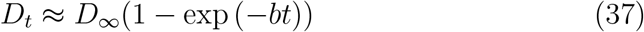

This is an author extrapolated expression, validated with curve fitting by varying the parameter *b*, which is approximately 0.155 for all *P*_*A*_ = *P*_*B*_ when *A* = 1. Comparing the fitted values for *b* for different values of *R*, a close fit is given by a quadratic incorporating for *R* and *R*^2^ (Eureqa, v1.12.1). The form of an expression for *b* can be given by

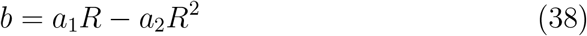

Again, *a*_1_ and *a*_2_ are constants but their fitted values for *P*_*A*_ = *P*_*B*_ are approximately *a*_1_ = 0.45 and *a*_2_ = 0.3 when *A* = 1. So *b ≈* 0.15(*R* + 2*R*(1*− R*)). The maximum rate of change in linkage disequilibrium occurs for unlinked loci and no change occurs when the loci are linked. All other values of *A* have similar expressions though the constants will differ. In general, for a given value of *R, b* is larger for lower values of *A* since their lower values of *D*_*∞*_ are reached more rapidly.

### 9.2. *Effects of recombination under disassortative mating where P*_A_ = 1 *− P*_*B*_

The effect of recombination in disassortative mating when *P*_*A*_ = 1 *− P*_*B*_ is the similar to the previous case. The main difference is the more rapid convergence to equilibrium in disassortative mating makes the constant *b* larger, closer to 1 for *A* = 1. Likewise the expression for *b* is *b ≈ R* + 2*R*^2^. Therefore the rate of recombination again increases the rate of convergence to equilibrium and linkage disequilibrium does not accumulate when the loci are completely linked.

### 9.3. *Effects of recombination under assortative mating where P*_*A*_ = 1 *− P*_*B*_

Recombination’s effect on convergence remains similar when *P*_*A*_ = 1 *− P*_*B*_, however, the dependence of *b* on *R*^2^ becomes much weaker as the magnitude of difference between |*P*_*A*_ *− P*_*B*_| gets larger so that it is nearly a linear dependence on *R*.

## 10. Identity Disequilibrium and Composite Linkage Disequilibrium

A key question addressed earlier is how the inbreeding and linkage disequilibrium effects determine the frequency of two-locus genotypes. In typical analyses of inbreeding, the two-locus genotype frequency is altered by identity disequilibrium which increases the frequency of double homozygotes and double heterozygotes and decreases the frequency of homozygote-heterozygotes to an extent greater than expected by just accounting for the effects of inbreeding a each locus alone.

In pure inbreeding problems, the identity disequilibrium variable effects are roughly symmetric and of equal magnitude for all two-locus genotypes. However, the identity disequilibrium is a direct result of the pedigree and its inbreeding and closely connected to the idea of the probability that both loci have two alleles that are identical by descent. In assortative mating problems though, while inbreeding like effects occur with the reduction of heterozygosity, these effects can be assumed to happen even in an infinite population with no drift effects where consanguineous matings are negligible. In addition, given some double homozygotes increase in frequency in most scenarios, such as *AABB* and *aabb*, while the other double homozygotes *AAbb* and *aaBB* as well as double heterozygotes decrease, it is questionable if the exact conceptualization of identity disequilibrium can be fully transferred to the assortative mating case.

Despite this, there are similar effects such as correlations in homozygosity across loci, even in the first generation where there is no linkage disequilibrium. Subsequent to the first generation of assortative mating, linkage disequilibrium clouds the picture so that there is an overall correlation in homozygosity but the separate contributions from effects deriving from identity disequilibrium and those of linkage disequilibrium are not clear nor easy to disentangle.

However, as explained in the section on recombination, when *R* = 0 there is no linkage disequilibrium and therefore the identity disequilibrium can be clearly analyzed without confounding.

Below we will show some results with the clearest being from the *P*_*A*_ = *P*_*B*_ case.

### 10.1. *Identity disequilibrium when P*_*A*_ = *P*_*B*_ *and R* = 0

When *R* = 0, there is no increase in linkage disequilibrium, but it is found that two-locus genotype proportions do change though the effect is due to identity disequilibrium and not linkage disequilibrium. There is not correlation amongst gametes, however, there is correlation in homozygosity which changes the two-locus genotype frequencies accordingly. The identity disequilibria were calculated using variations of Equation 18. There is a clear pattern across all *P*_*A*_ = *P*_*B*_ and across values of *A*. In the basic case of *A* = 1, the identity disequilibria vary by genotype and are summarized below

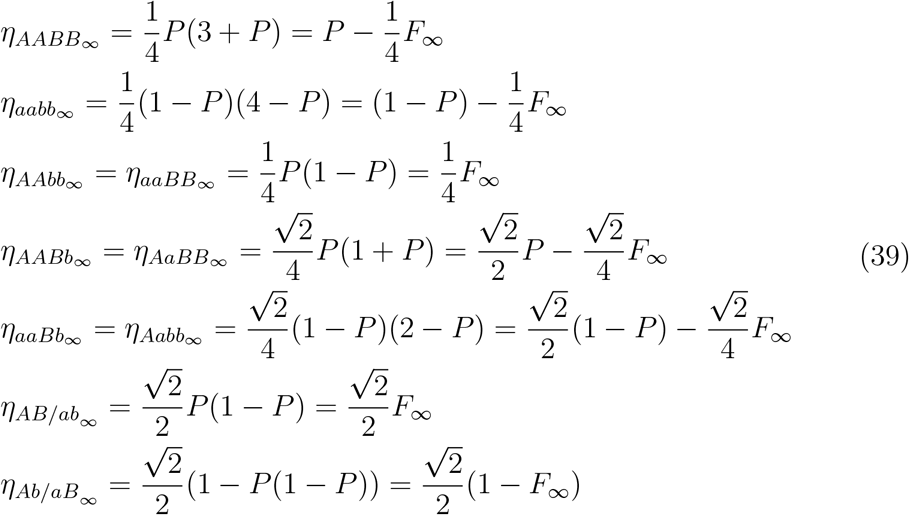

When *A <* 1 the basic relationships and equivalences between genotype frequencies that define identity disequilibria remain the same, only the variable coefficients change in value. A closed form solution for the changes of the coefficients for all values of *A*, however, was not entirely clear.

### 10.2. *Identity disequilibrium when P*_*A*_ = 1 *− P*_*B*_ *and R* = 0

Identity disequilibria in this case also follow qualitatively similar relationships but a closed form expression for the coefficients, even at *A* = 1 was not entirely clear. However, there are equivalency relationships between several of the identity disequilibria in this case.

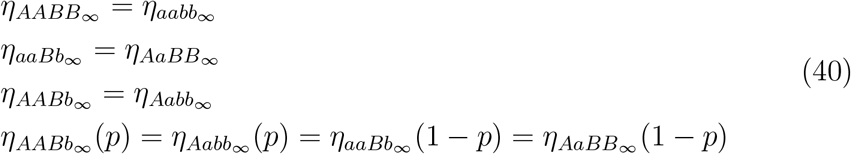

*<h2>10*.*3. Two-locus genotypes when P*_*A*_ = *P*_*B*_, *A* = 1 *and R >* 0

When *R >* 0 and *A* = 1, the equilibrium two-locus genotype frequencies were given in (Ghai, 1973). When *P*_*A*_ = *P*_*B*_ = *P* these can be given according to

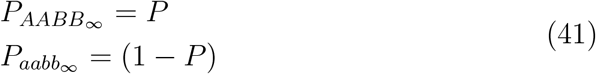

When *P*_*A*_ ≠ *P*_*B*_ there is a third double homozygous genotype present. When *P*_*A*_ *> P*_*B*_ the equilibrium genotype frequencies are

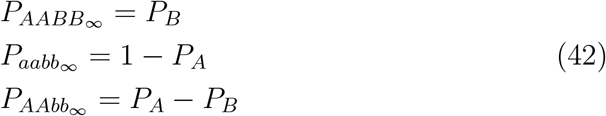

When *P*_*A*_ *< P*_*B*_ the equilibrium genotype frequencies are

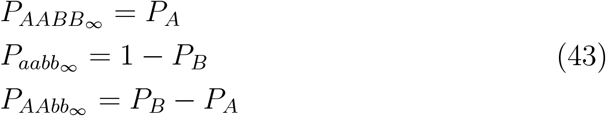

### 10.4. Composite linkage disequilibrium

Composite linkage disequilibrium, the measure of *P*_*AB/ab*_–*P*_*Ab/aB*_ is nearly zero in this context and also in the examples of previous sections. The two double heterozygotes tend to have the same equilibrium frequencies across all assumptions and thus compound linkage disequilibrium reflects this and does not seem to be an important or distinguishing metric in assortative mating.

## 11. Potential genetic assortative mating evidence in human populations

The foregoing theoretical work has established a basis for the overall effects of assortative mating on heterozygosity and linkage disequilibrium by finding exact or approximate solutions for the simple cases of *P*_*A*_ = *P*_*B*_ and *P*_*A*_ = 1 *− P*_*B*_. The strength of the general model, however, is that it can be used to analyze various situations which may not be amenable to closed form solutions but nevertheless accurately reflect biological reality.

There have been many works looking at assortative mating in human populations, primarily from phenotypic correlations amongst partners, but more recently using large amounts of SNPs and models to infer the genetic impact of assortative mating on complex traits (Robinson et. al., 2017; Yengo et. al., 2018). The effects of assortative mating on quantitative characters are similar (larger genetic variance, increased homozygosity). There are also similar challenges in that it can be difficult to disentangle genuine assortative mating for a phenotype from assortative mating based on a correlated character, population or social stratification that can make mates more similar by chance, and shared environmental influences. While two-locus traits have these same issues, the usually direct and binary effect of Mendelian traits makes the observation of genetic evidence such as homozygosity and linkage disequilibrium less ambiguous. There can be exceptions such as partial penetrance for a phenotype given a genotype or other evolutionary forces that can produce similar evidence as assortative mating. However, we will note these possible confounding effects, and eliminate those that are most infeasible, rather than present a model that claims to control for all possible explanations.

We will investigate two areas of likely phenotypic assortative mating using variations of the general model and population genetic data for non-Finnish Europeans from the 1000 Genomes (1000 Genomes Project Consortium, 2015) and Gnomad (Chen et. al., 2022) public databases to estimate correlations between phenotypes that most accurately represent the data. The non-Finnish European populations are selected due to the larger size of the dataset, more past studies analyzing assortative mating, as well as the fact that it has no recent historical population bottlenecks which may have non-assortative mating induced effects on homozygosity and linkage disequilibria.

### 11.1. Assortative mating amongst the deaf

Assortative mating amongst the deaf was one of the earliest examples debated when the science of heredity began in the nineteenth century. Decades before assortative mating was tackled by Jennings, Robbins, and Wright, Alexander Graham Bell among others raised an alarm about assortative mating between deaf individuals due to the rise of many schools and other institutions that supported them and facilitated partnering between deaf individuals (Bell, 1884). It was claimed that this assortative mating would create a sub-population of deaf individuals and increase the prevalence of hereditary deafness. Bell’s Volta Bureau funded a study by Edward Fay (Fay, 1898) that concluded that deaf marriages had not significantly raised the prevalence of deafness in America. In fact, then as now, many deaf parents have children without hearing impairment.

Bell and his successors did not know assortative mating alone does not change allele frequencies since the problem had not been well-defined, much less worked out. However, assortative mating can increase disease incidence through a rise in homozygosity. This argument was touched on in (Crow & Kimura, 1970) in their discussion on assortative mating though they believed the overall impact on the prevalence of deafness would be small unless the matings were consanguineous. Assortative mating in the context of cultural transmission of sign language was investigated in (Aoki & Feldman, 1991). Debate about the impact of assortative mating amongst the deaf has continued to this day. Some authors assert that the combination of assortative mating and relaxed selection against the deaf due to supportive institutions and the introduction of sign language can drive allele frequency increases of deleterious alleles in the population (Nance et. al., 2000; Nance & Kearsey, 2004; Blanton et. al., 2010). Others have disputed the selection effect and asserted the overall impact of assortative mating alone is weak (Braun et. al., 2020).

There is ample evidence for assortative mating between the deaf from survey data. Fay’s studies were analyzed in (Rose, 1975) to show that roughly 75% of profound deaf adults married other deaf adults. The numbers reported were even higher in the 1974 National Survey of the Deaf and ranged between 80-90% (Schein & Delk, 1974). More recent work in (Blanton et. al., 2010) showed an assortative mating percentage at 79% in alumni of Gallaudet University.

While there are many causes for deafness, with both environmental and biological factors, about 50% of pre-lingual hearing loss in developed countries is believed to be due to genetic causes and 70% of genetic hearing loss is classified as non-syndromic hearing loss. This means non-syndromic hearing loss accounts for 35% of all pre-lingual hearing loss (Putcha et.al., 2007). Non-syndromic hearing loss is differentiated from other hereditary forms of hearing loss in that it is hearing loss not associated with another broader syndrome such as Usher’s Syndrome. Hearing loss is typically the only effect of the disease with non-syndromic hearing loss. Non-syndromic hearing loss has a very diverse genetic etiology spanning many loci across multiple genes in both autosomal and sex chromosomes with alleles exhibiting recessive or dominant character. However, about 80% of the known variants that cause non-syndromic hearing loss are autosomal recessive (Putcha et.al., 2007).

The highest frequency variants causing non-syndromic hearing loss vary across global populations. Variants affecting the protein connexin 26 and associated with the gene *GJB2* account for 50% of the known autosomal recessive variants. There are two autosomal recessive variants which are most common in European descended populations: a frameshift variant caused by a G deletion, *35delG* (GrCH 38: 13-20189546-AC-A/rsID: rs80338939) which accounted for 60% of *GJB2* autosomal recessive variants in White cases a large US based study (Putcha et.al., 2007) and the missense substitution *101T >C* (also known as *M34T*) (GrCH 38: 13-20189481-A-G/rsID: rs35887622) which accounted for 12% of White cases in the same study. These are the most common variants in the US White and Hispanic patients in the study but are not the most frequent variants for African-American, Asian, or Ashkenazi Jewish patients though *GJB2* is still the most prominent affected gene across populations.

Complicating the genetic etiology, however, is the fact that there is frequently incomplete penetrance for the severe hearing loss phenotype amongst homozygous individuals with many of the variants. While frameshift variants such as *35delG* and *167delT* (GrCH 38: 13-20189414-CA-C/rsID: rs80338942), the most common variant in the Ashkenazi Jewish population, have a high penetrance of the severe deafness phenotype, SNP variants such as *101T >C* that cause missense changes are typically lower penetrance (Snoeckx et. al., 2005). In (Snoeckx et. al., 2005) 90% of genotyped individuals homozygous for frameshift variants had severe to profound deafness compared to only 21% of individuals homozygous for missense variants. This leads to the interesting situation where the allele frequency of the *101T >C* variant in non-Finnish European populations is 0.014 compared to *35delG* which is only 0.008 according to the Gnomad 4.0.0 data. However, there usually is a higher penetrance of severe deafness phenotypes in *35delG* homozygotes.

The most favorable situation to investigate would be using two frameshift variants with high penetrance to investigate the effects of assortative mating at two loci. However, the second highest frequency frameshift variant, *167delT*, is at significantly lower frequency in the non-Finnish European population in Gnomad v4.0.0, having zero homozygotes, and only has three homozygotes in the Ashkenazi Jewish population. Likewise *35delG* has no homozygotes in the Ashkenazi Jewish population in Gnomad. Therefore we will investigate if there is evidence of assortative mating at either the single locus of *35delG* or *101T >C* or jointly with *35delG* and *101T >C* in the non-Finnish European population.

Given the separation of the two loci by less than 100 bp, they are completely linked and linkage disequilibrium is not expected to be generated by assortative mating. Therefore, the main focus to analyze possible assortative mating is an excess of homozygosity at both loci. The data on the allele frequencies, genotype frequencies, and Hardy-Weinberg chi-square test p-values for *35delG* and *101T >C* in non-Finnish European populations and *167delT* in Ashkenazi Jewish populations based on Gnomad v4.0.0 are in Table 5.

**Table 5:**
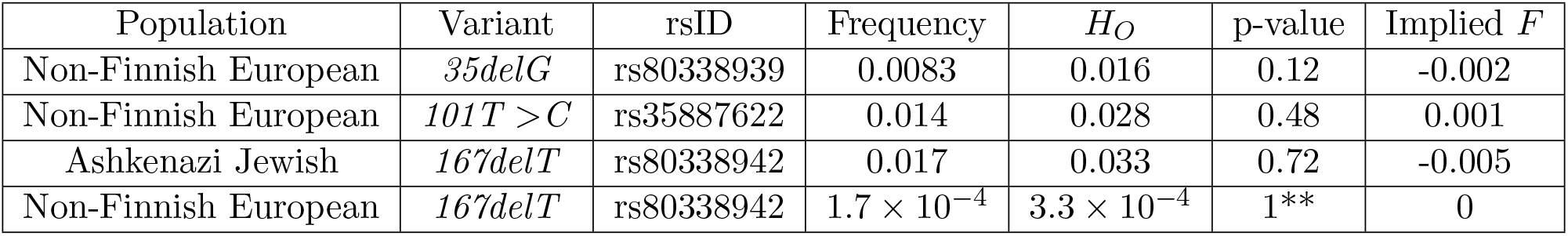
Populations and genetic data for common autosomal recessive variants that can lead to hearing impairment. The observed heterozygote frequency is designated by *H*_*O*_ Data are based on allele and homozygote frequency from Gnomad v4.0.0 and p-values and *F* are calculated with the R package HardyWeinberg (Graffelman, 2015).^**^Used Hardy-Weinberg Exact Test due to low homozygote count.

Contrary to expectations, none of the variants reach a significant p-value threshold that would suggest deviation from Hardy-Weinberg equilibrium, and the estimated values of *F*, are possibly due to chance. Average values of *F* at loci in European populations typically range anywhere from near 0 up to 0.004 (Pemberton & Rosenberg, 2014) which can obscure the ultimate cause of the value of *F* in this case. This is a surprising result since there is clear survey evidence of assortative mating amongst the deaf. What may be happening is that the results of such assortative mating are muted due to the low penetrance of severe deafness for many genotypes. This would make assortative mating weaker since individuals who are homozygous but not impaired would likely have children with the general population. Linkage disequilibrium data shows linkage equilibrium for the phase date in Gnomad v2.1.1 and near zero *r*^2^ in 1000 Genomes as well.

However, given the relatively high penetrance of frameshift mutations *35delG* and *167delT*, the lack of significant excess homozygosity may indicate that the assortative mating is too weak at a population level to have a significant impact.

To investigate the results of assortative mating on homozygosity in the population, a recursive model was developed that separated all ten two-locus genotypes into two phenotype groups. A group where the genotypes would indicate autosomal recessive non-syndromic deafness and one with normal hearing where all genotypes are carriers or do not have either of the two common *GJB2* disease variants. Where the *35delG* locus is *A/a* and the *101T >C* locus is *B/b*, the deaf phenotype group consists of the clear homozygotes of *35delG aaBB, aaBb*, and *aabb* as well as the clear homozygotes of *101T >C AAbb* and *Aabb*. All other genotypes belong to the non-impaired phenotype group. The recombination rate between loci was set to *R* = 0. The simulation was run for 20 generations or the approximate time for several centuries of assortative mating.

Using the above defined phenotype groups and Gnomad v4.0.0 allele frequencies, the model was recursively run and the expected equilibrium values of *F* for each locus were plotted. The results in Figure 8 show that even under strong assortative mating and assuming full penetrance of the genotypes, the fixation index does not become very high. When *A* = 1, the *F* values range from 0.015 to 0.026. Lower values of *A* yield fixation indices less than 0.005 consistent with results from (Braun et. al., 2020).

**Figure 8.**
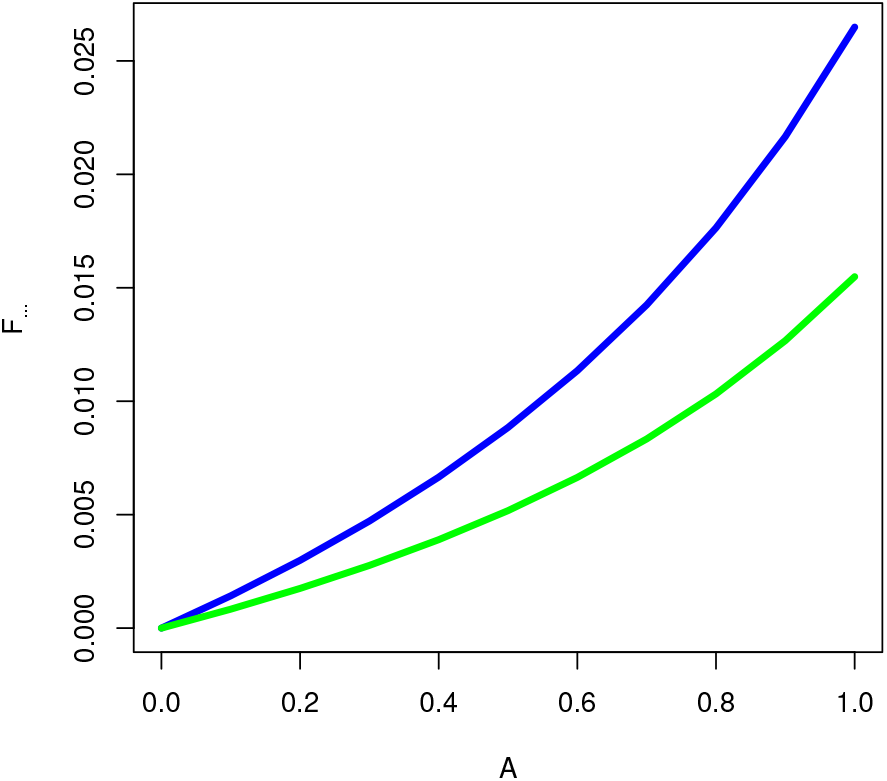
A plot of the values of *F*_*∞*_ vs. *A* for the two most frequent loci causing non-syndromic deafness in European populations. The blue line represents the fixation index for *35delG* and the green line represents the fixation index for *101T >C*.

One could argue that a single locus assortative mating model is more cogent. For a locus with complete dominance and only two phenotypes, equation 4.6.7 in (Crow & Kimura, 1970) expresses the relationship between *F, q* the recessive allele frequency, and *m* as

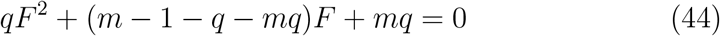

However, given the negative fixation index for the most common causal variant, *35delG* in both Gnomad v4.0.0 as well as v2.1.1, a single locus model would still imply no or slightly disassortative mating at that locus. For *101T >C*, even assuming full penetrance of the homozygous recessive genotype, a value of *F*_*∞*_ of 0.001 would imply a *m ≈* 0.07. One interesting point is that in the smaller Gnomad v2.1.1 dataset, the frequency of *101T >C* is similar at 0.013 but the measured *F* is 0.009. Using similar equations and arguments for single locus assortative mating, *m ≈* 0.42. However, conclusively assigning this value for *F* to assortative mating still encounters the issue that the higher penetrance and more common deafness causing variant *35delG* still has a negative *F* of -0.003 in v2.1.1 which would not be the expected outcome in any assortative mating scenario.

Given these results and the available genetic data it seems reasonable to conclude that while assortative mating is definitely occurring amongst the deaf, its impact as an independent factor on the population genotype frequencies of homozygous genotypes that have a high propensity for deafness is relatively mild. The largest drawback to this data, however, is the European population data is not limited to the United States. If assortative mating is much weaker in other European descended populations, this could mask the effect of assortative mating in the US. Finally, the hypotheses of increased fertility leading to a higher frequency for deleterious alleles leading to deafness cannot be adequately evaluated using this paper’s framework alone.

### 11.2. Assortative mating based on eye color

Many well-known physical phenotypes such as height, skin color, or hair texture have a polygenic character. Eye color is one, however, that is determined in the majority of cases by a relatively small number of known loci, some of which interact in an epistatic fashion. One example is blue eye color which is common in, but not unique to, European descended populations. There are many loci that influence eye color, however, for blue eyes three loci play a prominent role in prediction as outlined in genetic studies as well as eye color forensic prediction tools such as IrisPlex (Eiberg et. al., 2008; Mengel-From et. al., 2010; Walsh et. al., 2011).

The primary and necessary locus to generate blue eyes is an intronic variant on HERC2 gene that regulates expression on the nearby OCA2 gene (rsID: rs12913832/GrCH 38:15-28120472-A-G) (White & Rabago-Smith, 2011). This autosomal recessive variant affects the regulation of expression for OCA2 that can lead to blue eyes with high penetrance when there is homozygosity of GG. Homozygotes for the dominant allele and heterozygotes usually, though not always, have brown eyes. In many global populations, absent admixture, the minor allele G is at a low frequency or even does not exist and blue eyes are absent. However, further influence on the likelihood and color of blue eyes is given by two additional autosomal recessive loci, (rsID:rs12896399 / GrCH 38:14-92307319-G-T) and (rsID: rs16891982 / GrCH 38: 5-33951588-C-G). Having homozygosity for the minor allele at either or both of these loci increases the tint of blue eyes in an individual.

Past research indicates that many studies have found that couples practice assortative mating with respect to eye color. The earliest work on such assortative mating was by Karl Pearson who used data collected by Francis Galton to find correlation between spouses on many attributes such as stature and eye color, which was measured by a non-continuous range of tints (Pearson, 1900). In two different calculations, Karl Pearson measured the correlation between mates for eye color. When he used 8 different categories in (Pearson, 1900) he found *m* = 0.10 ± 0.04. Using five categories in (Pearson, 1907), by combining some of the less represented tints with more frequent ones of a similar color, *m* = 0.26 ± 0.03. Later work by (Elston, 1963) on couples in Uppsala, Sweden found evidence of assortative mating for blue and dark eye color. Count tables indicate that the correlation for couples with blue eyes is *m* = 0.17 (calculations by author from the Table 2 in (Beckman, 1962)). However, in (Beckman, 1962) the author’s separate work on Lapp’s in Sweden failed to find convincing evidence for assortative mating. In this measurements though, the author used different eye color categorizations (light or mixed vs. brown eyes) and the author also acknowledged the high frequency of brown eyes in the population could make assortative mating much more difficult. In (Laeng et. al., 2007), experiments using volunteer evaluations of faces as well as surveys of the eye color of male and female participants and their partners gave some evidence for assortative mating between men and women with blue eyes with less evidence for that between those with brown eyes. This author’s calculations based off of Tables 1 and 2 indicate correlations for blue × blue eyes at *m* = 0.35 in the male participant group and *m* = 0.13 in the female participant group.

Here, we will use population genetic data as well as simulations to estimate the assortative mating that excess homozygosity and linkage disequilibrium suggest. It is important to note that it cannot be clear from this data if the assortative mating is directly caused by solely eye color or a confounding variable such as sub-population structure or correlated phenotypes like skin pigmentation that may produce a similar effect.

One advantage of studying the three loci determining blue eye color is their presence on different chromosomes means they are completely unlinked and can generate linkage disequilibrium due to assortative mating. Therefore, unlike the previous example we will use not just single locus homozygosity but also linkage disequilibrium and two-locus genotype frequencies. However, since Gnomad v2.1.1 only provides phase data for loci on the same gene, we will rely on the 1000 Genomes European population data (excluding the Finnish participants) to measure linkage disequilibrium and two-locus genotypes.

In Table 6 are the allele and genotype frequencies as well as p-values from the Chi-Square test for Hardy-Weinberg equilibrium and the implied value of *F* for data in 1000 Genomes Phase 3 as well as Gnomad v.4.0.0. From the Gnomad data perspective, each locus is not in Hardy-Weinberg equilibrium and have substantial values for the fixation index, larger in the 1000 Genomes data set. In Table 8, is a linkage disequilibrium matrix based on the 1000 Genomes data. Despite being on different chromosomes, there is sizable linkage disequilibrium between variants rs12913832 and rs16891982. The two-locus genotype frequencies (See Table 7) also are substantially different than the equilibrium expectation where the two-locus double homozygote genotype for the blue -eye associated variants at rs12913832 and rs16891982 has a frequency of 0.34 compared to the equilibrium expectation of 0.28.

**Table 6:** Population genetic data on the main loci determining blue eye color. Data are based on allele and homozygote frequency from 1000 Genomes and Gnomad v4.0.0 and calculated with the R package HardyWeinberg (Graffelman, 2015). 1000 Genomes data Phase 3 with data from all European (EUR) participants except Finnish participants (FIN). Gnomad data uses all data for non-Finnish Europeans. ^**^Used Hardy-Weinberg Exact Test due to low homozygote count.

| Database | Variant rsID | Frequency | $H_O$ | p-value | Implied $F$ |
| --- | --- | --- | --- | --- | --- |
| 1000 Genomes | rs12913832 | 0.57 | 0.42 | 0 | 0.15 |
| 1000 Genomes | rs12896399 | 0.58 | 0.46 | 0.22 | 0.06 |
| 1000 Genomes | rs16891982 | 0.93 | 0.11 | 0.09** | 0.09 |
| Gnomad v4.0.0 | rs12913832 | 0.76 | 0.34 | 0 | 0.06 |
| Gnomad v4.0.0 | rs12896399 | 0.44 | 0.49 | 0.03 | 0.01 |
| Gnomad v4.0.0 | rs16891982 | 0.97 | 0.06 | 0 | 0.03 |

**Table 7:** Actual and (expected) frequencies for two-locus genotypes of rs12913832 and rs16891982. Allele symbols are *A* for rs12913832 and *B* for rs16891982.

| | $BB$ | $Bb$ | $bb$ |
| --- | --- | --- | --- |
| $AA$ | 0.34 (0.28) | 0.02 (0.04) | 0 |
| $Aa$ | 0.36 (0.43) | 0.05 (0.06) | 0 |
| $aa$ | 0.18 (0.16) | 0.04 (0.02) | 0 |

**Table 8:** *D*^*′*^ values between causative loci for blue eye color using 1000 Genomes EUR population minus the Finnish (FIN) participants.

|  | rs12913832 | rs12896399 | rs16891982 |
| --- | --- | --- | --- |
| rs12913832 | * | 0.02 | 0.25 |
| rs12896399 |  | * | 0.01 |
| rs16891982 |  |  | * |

Therefore, there is at least data that suggests that population genetic effects due to assortative mating are possible. The excess homozygosity at each locus as well as the increased frequency of the double homozygote are important since other forces such as selection and admixture can create linkage disequilibrium but do not lead to excess homozygosity in most cases. Drift can create both excess homozygosity and linkage disequilibrium, for example in a population bottleneck, however, the linkage disequilibrium created is strongly dependent on linkage between the two loci and exponential population growth, such as European populations have experienced for many centuries now, rapidly reduces the linkage disequilibrium making it shortlived (Slatkin, 1994). Likewise, consanguineous mating can lead to excess homozygosity but usually not substantial linkage disequilibrium and the values of *F* shown are substantially higher than to be expected by inbreeding in these populations as mentioned previously (Pemberton & Rosenberg, 2014). Since only the 1000 Genomes data set has two-locus and linkage disequilibrium data, we will look at two models of the recursive equations to try to reproduce the results. Though the alleles that generate blue eyes act in a recessive fashion, they are the major alleles and will be designated as *A* for rs12913832 and *B* for rs16891982. The first model has two phenotype groups: *AABB* or blue eyes alone in one phenotype group and all other genotypes in the second phenotype group. The second model uses three phenotype groups: one for *AABB* (blue eyes), one for *AaBB* (usually brown eyes) and the rest in the third phenotype group. This second model was proposed by analyzing the two-locus genotype frequency data in 1000 Genomes where *AABB* and *AaBB* have the largest frequency differences from the equilibrium expectation frequency (Table 7). In both models, the recombination rate is set to *R* = 1*/*2 and the simulation run for 30 generations where the equilibrium value was approximated in all cases.

For the 1000 Genomes data the simulation used both models and determined the value of *A*, in increments of 0.01, that produced the minimum sum of square difference between the simulation frequencies and measured frequencies for *AABB* and *AaBB*. For the Gnomad v4.0.0 data the value of *A* that minimized the sum of square difference between each frequency of heterozygosity from the simulation and the measured values was used to select the closest value for *A*. The results are shown in Table 9. The second model with three phenotype groups produced the closest results for both data sets.

**Table 9:** Simulation results for assortative mating for two-locus genotypes underlying the phenotype for blue eyes showing the best fit value for *A* given a phenotype group model and population genetic data from one of two datasets. The variants are rs12913832 (*A/a*) and rs16891982 (*B/b*) in Non-Finnish European populations from either the 1000 Genomes or Gnomad v4.0.0 datasets. Model I indicates two phenotype groups with group 0 containing *AABB* and group 1 all the other genotypes. Model II indicates three phenotype groups with group 0 containing *AABB*, group 1 containing *AaBB* and group 2 all the other genotypes.

|  | Population Data |  |  |  |  |  |  |
| --- | --- | --- | --- | --- | --- | --- | --- |
| Dataset | $pA$ | $pB$ | $F_A$ | $F_B$ | $D'$ | $P_{AABB}$ | $P_{AaBB}$ |
| 1000 Genomes | 0.57 | 0.93 | 0.15 | 0.09 | 0.25 | 0.34 | 0.36 |
| Gnomad v4.0.0 | 0.76 | 0.97 | 0.06 | 0.03 | N/A | N/A | N/A |

Table 9: Simulation results for assortative mating for two-locus genotypes underlying the phenotype for blue eyes showing the best fit value for $A$ given a phenotype group model and population genetic data from one of two datasets.
|  |  | Simulation Data |  |  |  |  |  |
| --- | --- | --- | --- | --- | --- | --- | --- |
| Dataset | Model | $m$ | $F_A$ | $F_B$ | $D'$ | $P_{AABB}$ | $P_{AaBB}$ |
| 1000 Genomes | Model I | 0.46 | 0.178 | 0.018 | 0.18 | 0.34 | 0.34 |
| 1000 Genomes | Model II | 0.35 | 0.16 | 0.06 | 0.24 | 0.34 | 0.35 |
| Gnomad v4.0.0 | Model I | 0.14 | 0.056 | 0.005 | 0.06 | 0.56 | 0.32 |
| Gnomad v4.0.0 | Model II | 0.14 | 0.06 | 0.03 | 0.08 | 0.56 | 0.32 |

In particular, for the 1000 Genomes data, the closest value is *A* = 0.35 which matches *P*_*AABB*_ = 0.34 and *P*_*AaBB*_ = 0.35 and gives a linkage disequilibrium value of *D*^*′*^ = 0.23, while *F*_*A*_ = 0.16 and *F*_*B*_ = 0.06. For the Gnomad data, the value of *A* is smaller with a best fit for *A* = 0.14 with *F*_*A*_ = 0.06 and *F*_*B*_ = 0.03. Unfortunately two-locus and linkage disequilibrium data is unavailable with Gnomad for additional validation.

These results seem to suggest that genotype frequencies and linkage disequilibrium in the European population support the case for assortative mating due to eye color. In particular, since there is only one genotype in the two main phenotypes groups, *A* = *m* for their assortative mating. The closest fit of *A* = 0.14 in the larger Gnomad v4.0.0 dataset closely matches past correlations by (Pearson, 1900) and (Elston, 1963) as well as the female participant correlation calculated from (Laeng et. al., 2007). The 1000 Genomes estimate of *A* = 0.35 is higher than any observational estimate and may be an overestimation due to some aspect of the sample. However, this genetic data cannot rule out the possibility this effect is not by eye color per se but a close proxy. One such proxy could be a strongly correlated phenotype, due to linkage disequilibrium with the eye color variants or pleiotropy. Another could be a sub-population with a higher frequency for the blue eye alleles. Also, while this analysis makes assortative mating a plausible cause of the genetic data observed, it cannot conclusively eliminate other explanations.

## 12. Discussion

The general effects of assortative mating on population genotype frequencies have long been known. However, descriptive solutions for equilibrium values of population genetic variables where two loci underlie a phenotype have rarely been given outside of the narrow conditions of *P*_*A*_ = *P*_*B*_ = 1*/*2 and *m* = 1. Much of this was due to the complex nature of recursive assortative mating calculations and the lack of an analytical framework for most conditions. This paper was written to solve most of these issues. First, by introducing a clear methodology to recursively iterate a population without agent modeling or simulation and second adopting this to symbolic algebra systems that can produce interpretable output that can help in deriving exact solutions.

While exact, closed form solutions were difficult to find outside of assortative mating for *P*_*A*_ = *P*_*B*_, the first general expressions for linkage disequilibrium were derived and close approximations for the fixation index were derived for assortative and disassortative mating where *P*_*A*_ = 1 *− P*_*B*_. In addition, several valuable insights were discovered or confirmed including the facts that recombination slows the approach to the equilibrium values for variables but not change the absolute equilibrium values, that assortative mating does not generate linkage disequilibrium between completely linked loci, and that a form of identity disequilibrium plays a role in determining two-locus genotype frequencies.

These general insights as well as the overall model can be applied to assortative mating in natural populations. Where assortative mating phenotype groups and correlations are known, the model can help predict the expected genotype frequencies and linkage disequilibrium in a population. When only population genetic data is available, the model can be used to find both likely phenotype groups as well as the assortative mating correlation that best fits the genetic data. The flexibility of being able to approach assortative mating problems from both perspectives will hopefully improve understanding of the probable strength of assortative mating and its impact on populations in future investigations.

## 13. Data Availability

Basic code in Python used for the model simulations is available from the author on request.

Declarations of interest: none. This research did not receive any specific grant from funding agencies in the public, commercial, or not-for-profit sectors.

